# The RNA-binding proteins CELF1 and ELAVL1 cooperatively control RNA isoform production

**DOI:** 10.1101/373704

**Authors:** Géraldine David, David Reboutier, Stéphane Deschamps, Agnès Méreau, William Taylor, Sergi Padilla-Parra, Marc Tramier, Yann Audic, Luc Paillard

## Abstract

ELAVL1 and CELF1 are RNA-binding proteins that are involved in alternative splicing control. To explore their functional relationship, we looked for mRNAs that are differentially spliced following the depletion of CELF1, ELAVL1, or both. We found that these proteins control the usage of their target exons in the same direction. Double depletion has a greater effect than individual depletions, showing that CELF1 and ELAVL1 exert additive control. To confirm these results, we carried out RT-PCR on the alternative cassette exons of several mRNAs, including *CD44, WNK1, PHACTR2, MICAL3, SPTBN1*, and *PPP3CB*. Using FRET, we found that CELF1 and ELAVL1 directly interact in cell nuclei. We demonstrated that the combined levels of *CELF1* and *ELAVL1* are a valuable biomarker in several cancers, even when their individual levels may yield very limited information. *CD44* alternative splicing probably accounts in part for the effects of *CELF1* and *ELAVL1* levels on patient survival. These data point to strong functional interactions between CELF1 and ELAVL1 in the control of mRNA isoform production, resulting in significant impacts on human pathology.

## INTRODUCTION

Alternative splicing is the process allowing several different mRNA molecules to be synthesised from a single gene. Deep sequencing approaches in the last decade have revealed that virtually all human genes produce at least two alternative mRNA isoforms (Wang *et al*, 2008; Pan *et al*, 2008). The key signals for splicing are the branch point and the 5’ and 3’ splice sites. These are all strictly required for splicing, and their core sequences are highly conserved. Consequently, alternative splicing control is seldom exerted through these sites themselves. Cis-acting splicing regulatory elements (splice enhancers or silencers) are instead found in introns or exons, and they regulate sites that can be tens or even hundreds of nucleotides apart. Their combined effects define a splice “code” that determines which mRNAs are produced when a pre-mRNA molecule is processed in any given cell type and at any differentiation stage (Wang & Burge, 2008; Fu, 2004).

Essentially, splicing regulatory elements act by binding RNA-binding proteins (RBPs). However, pre-mRNAs are bound by multiple RBPs, and splicing outputs depend on the relationships between these RBPs. For example, QKI competes with PTBP1 to control the inclusion of the *CAPZB* exon 9 (Hall *et al*, 2013), while HNRNPLL and HNRNPL act as antagonists to control the splicing of *CHRNA1* mRNA (Rahman *et al*, 2013). Conversely, RBFOX1 stimulates HNRNPM-mediated control of the splicing of many exons (Damianov *et al*, 2016). Hence, looking at a splicing regulation exerted by a single RBP is insufficient for predicting splicing patterns, and there are not enough results from experiments in living cells done on a genome-wide scale to provide a global view of the functional interactions between RBP pairs (Dassi, 2017).

Here, we address the functional relationships of two RBPs involved in alternative splicing control: CELF1 (CUGBP Elav-like family member 1, also called CUGBP1), and ELAVL1 (ELAV-like RNA-binding protein 1, also known as HuR). Both proteins interact with their RNA ligands through three specific domains nammed “RNA-recognition motifs”. The first two domains share 31% identity between CELF1 and ELAVL1, and the third shares 46%. Both proteins control alternative splicing (Kalsotra *et al*, 2008; Philips *et al*, 1998; Tang *et al*, 2015; Chang *et al*, 2014), and they both oligomerise (Cosson *et al*, 2006; Fialcowitz-White *et al*, 2007). They also co-immunoprecipitate, indicating that they bind to the same RNAs (Le Tonquèze *et al*, 2010). Finally, the phenotypes caused in mice by disrupting their genes look much alike. In a pure genetic background, inactivating either *Elavl1* or *Celf1* causes embryo death (Cibois *et al*, 2012; Katsanou *et al*, 2009). Conditional inactivation of *Elavl1* (Chi *et al*, 2011) and constitutive inactivation of *Celf1* in a mixed genetic background (Boulanger *et al*, 2015) showed that both ELAVL1 and CELF1 are required for spermatogenesis.

Despite their resemblances, ELAVL1 and CELF1 are highly divergent. The human reference isoform for CELF1 is significantly longer than the ELAVL1 one (489 versus 326 amino acids). This is due to a longer linker region between the second and the third RNA recognition motifs, a difference that probably results in new properties. Most importantly, the RNA binding specificities of these two proteins differ: CELF1 interacts with GU-rich elements, while ELAVL1 interacts with AU-rich ones. CELF1 binding to GU-rich elements most often leads to mRNA decay and represses or stimulates translation (Cibois *et al*, 2010; Chaudhury *et al*, 2016; Vlasova *et al*, 2008; Masuda *et al*, 2012), whereas ELAVL1 binding to AU-rich elements generally stabilizes the bound mRNA (Mukherjee *et al*, 2011; García-Domínguez *et al*, 2011; Lafarga *et al*, 2009; Chen *et al*, 2002; Peng *et al*, 1998). Their limited similarities make this pair of proteins attractive for the exploration of RBP relationships in alternative splicing control. Here, we show that CELF1 and ELAVL1 control together the splicing of their target genes, and that they directly interact in cell nuclei. Finally, we demonstrate the prognostic value for various cancers of looking at the combined mRNA levels of *CELF1* and *ELAVL1*, indications that are unseen when the levels are considered separately. Our discoveries point to strong functional interactions between CELF1 and ELAVL1 that were not expected given their weak affinity and dissimilar molecular functions, thus underlining the clinical importance of simultaneously considering the molecular functions of RNA-binding proteins.

## RESULTS AND DISCUSSION

### CELF1 and ELAVL1 similarly control pre-mRNA splicing

We used RNA interference to deplete CELF1 and ELAVL1 either separately or simultaneously in HeLa cells. To reduce toxicity, we used siRNA quantities which left at least 25% of these proteins. The two proteins were knocked down equally with single and double siRNAs, and there was no apparent cross regulation between the proteins (Figure S1A). Having established our experimental conditions, we extracted RNAs from control and depleted cells to analyse RNA splicing by exon array hybridization. Exon arrays are used to infer splicing patterns, with results comparable to those from deep RNA sequencing (Zhang *et al*, 2015; Raghavachari *et al*, 2012). For each of the 117,841 probes, we calculated a normalized exon probe value corresponding to the exon probe’s fluorescence intensity normalized by the expression of the exon probe’s gene. Based on these values, replicate conditions cluster together (Figure S1B). We filtered the exon probes of interest according to two criteria: they should target a gene that is reasonably expressed (among the 75% most-expressed genes under at least one condition); and based on CLIP-seq experiments carried out in HeLa cells, they should target an RNA ligand of both CELF1 and ELAVL1 (Table S1) (Le Tonquèze *et al*, 2016; Uren *et al*, 2011). This resulted in 18,714 exon probes in 1,299 genes. Of these, 62 exon probes in 57 genes have a significantly different normalized value in at least one test (depletion) compared with the control condition (control siRNA-transfected cells), at a false discovery rate (FDR) of 0.1. Table S2 lists these probes with their splicing indices (SIs), the log-ratios of normalized exon values in depleted cells compared to the normalized exon values in control cells.

We used the MISO annotation to classify the 62 differential exon probes (Katz *et al*, 2010). The distributions of differential RNA processing events upon depletion of either or both proteins are similar (Figure 1A). Alternative last exons are the most enriched, but skipped exons also represent a large number of events. We explored whether CELF1 and ELAVL1 repress or stimulate the use of each of the 62 exons covered by the probes. Figure 1B shows that all except one of the probes repressed by ELAVL1 are also repressed by CELF1, and that all but three of the probes stimulated by ELAVL1 are also stimulated by CELF1. This reveals that CELF1 and ELAVL1 essentially control splicing patterns in the same direction (*p* = 2 × 10^−10^, chi-square test).

**Figure 1.**
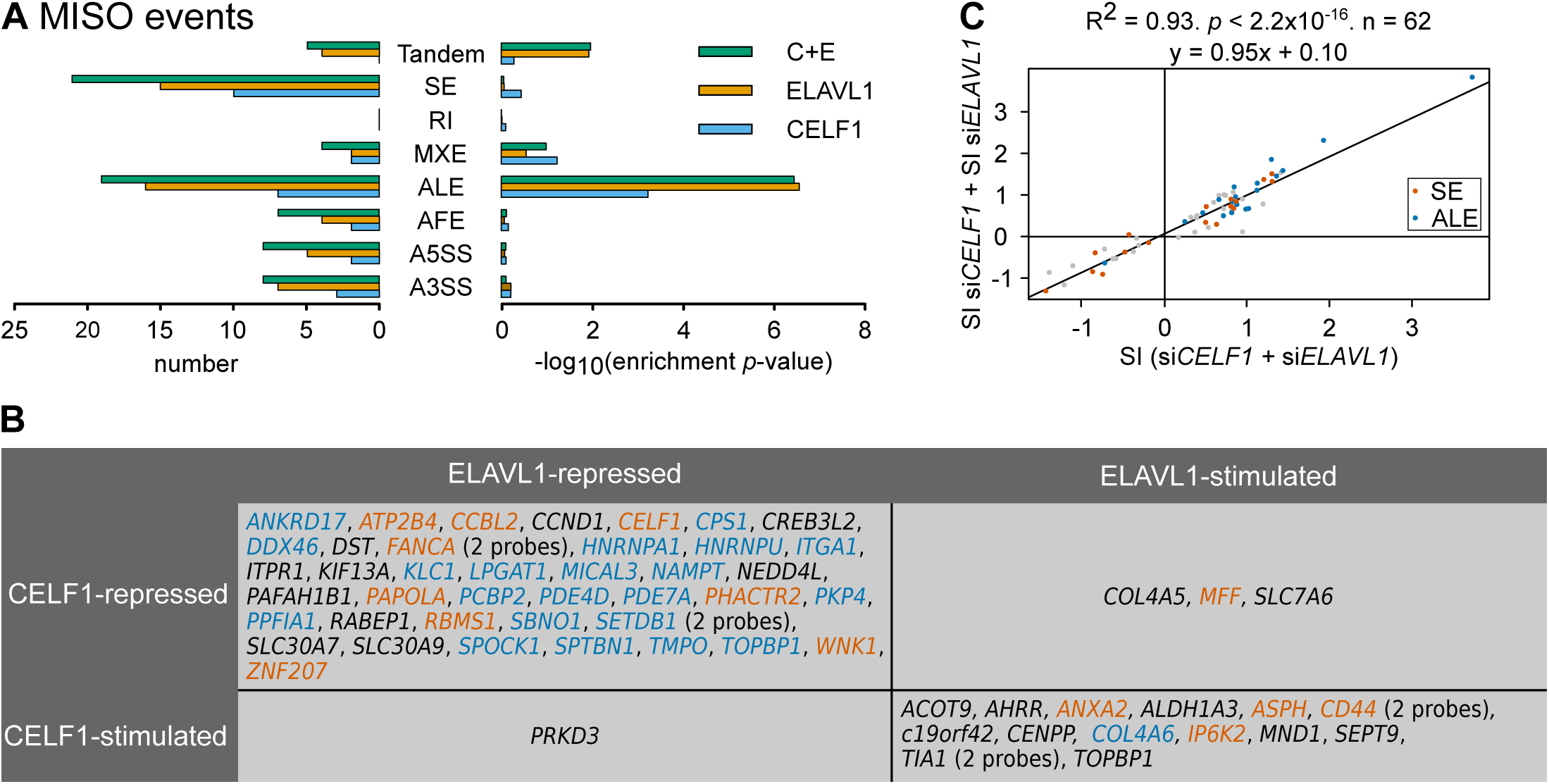
Alternative RNA isoform production in CELF1- and ELAVL1-depleted cells. **A**, *Left*, number of significantly differential probes affiliated with the indicated MISO RNA processing events after depleting CELF1 (light blue), ELAVL1 (orange), or both (green). *Right*, enrichment *p*-values of each MISO RNA processing event within the set of 62 differential probes. A3SS and A5SS, alternative 3’ and 5’ splice sites; AFE and ALE, alternative first and last exons; MXE, mutually exclusive exons; RI, retained introns; SE, skipped exons; Tandem, tandem untranslated region (alternative cleavage-polyadenylation sites). **B**, List of genes containing an exonic region that is either repressed (positive splicing index after depletion) or stimulated (negative splicing index) by CELF1 or ELAVL1. Alternative last exons and tandems are blue, and skipped exons are red. **C**, Sum of the splicing indices in CELF1- and ELAVL1-depleted cells plotted against the splicing indices in cells deprived of both proteins.

We wondered how the double depletion of CELF1 and ELAVL1 effects splicing. To explore this, we plotted the sum of the splicing indices after each single depletion against the splicing indices after double depletion (Figure 1C). These values are highly correlated (*R*^*2*^ = 0.93). Importantly, the slope of the regression line is close to 1 (95% confidence interval 0.88-1.02). Hence, the contribution of the two depletions to the splicing patterns in the doubly depleted cells equals the expectations from the additive effects of the individual depletions. This does not support any synergy nor redundancy between CELF1 and ELAVL1 in controlling RNA splicing patterns, and rather reveals additive controls.

### Validation of splicing regulation mediated by CELF1 and ELAVL1

We chose a cassette exon of *WNK1* to check the microarray results by RT-PCR. *WNK1* has a broad expression pattern. It encodes a kinase with a key role in kidney and neural cell functioning (Rodan & Jenny, 2017). It has a cassette exon that is regulated in a tissue-specific manner and which contributes to WNK1 protein level control by encoding a binding site for the E3 ubiquitin ligase NEDD4-2 (Vidal-Petiot et al, 2012; Roy et al, 2015). Our microarray data reveal that both CELF1 and ELAVL1 repress the usage of this cassette exon (Table S2). By RT-PCR, *WNK1* splicing pattern in non-transfected cells is very similar to that in cells transfected with either the control siRNA used in microarray experiments or an anti-luciferase siRNA (Figure 2A, lanes 1-3). This shows that the control siRNA is essentially neutral when it comes to splicing patterns. We also observed that the inclusion of the cassette exon is stimulated in a similar manner when using the siRNAs against *CELF1* together or separately (lanes 4-7), and the same is true for the siRNAs against *ELAVL1* (lanes 8-10). Therefore, the changes observed in splicing patterns must not be due to siRNA off-target effects.

**Figure 2.**
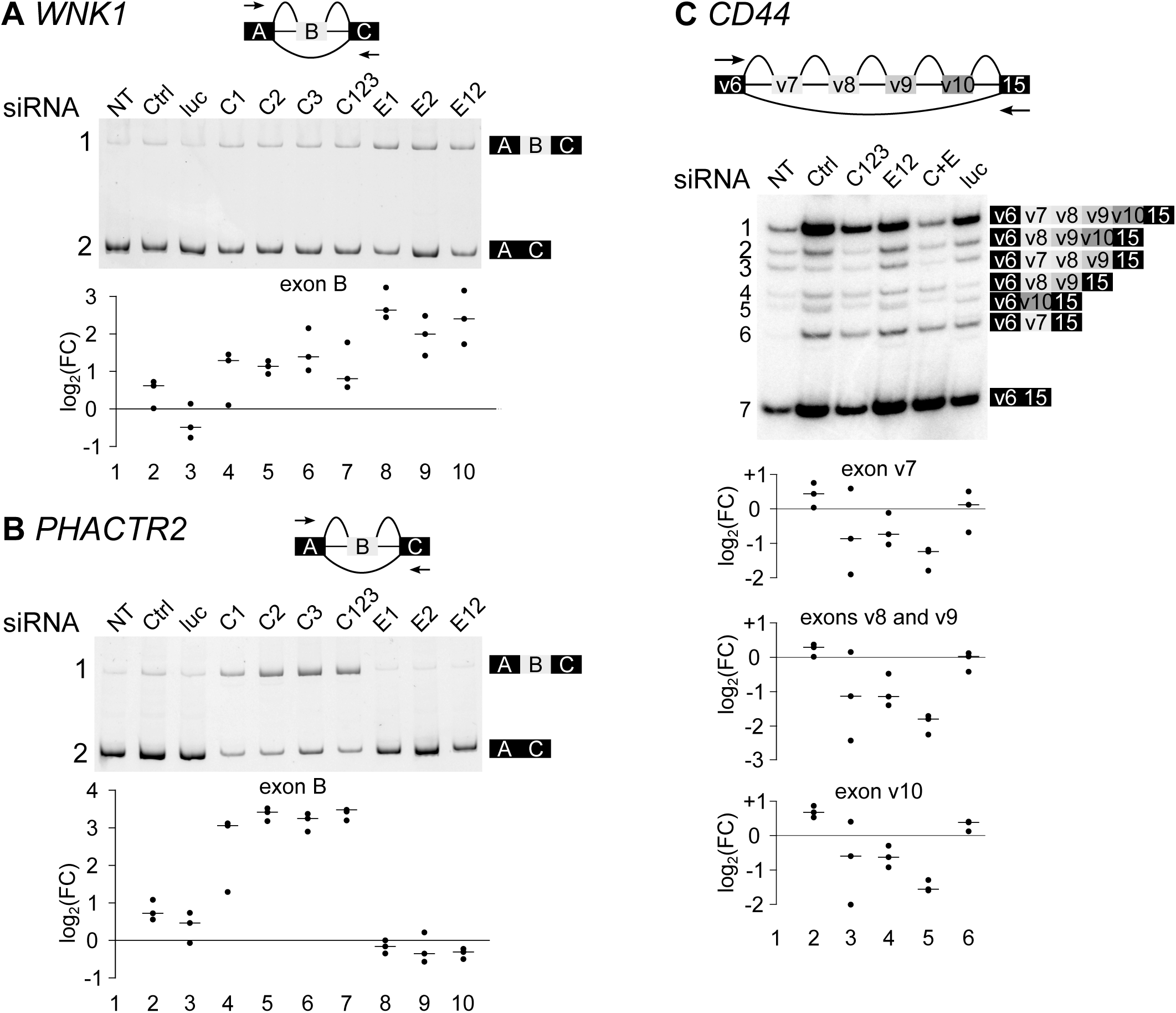
Examples of cassette exons controlled by CELF1 and ELAVL1. **A**, In *WNK1* pre-mRNA, a cassette exon labelled B is repressed by CELF1 and ELAVL1 in microarray experiments. We carried out RT-PCR with primers located in the flanking exons labelled A and C (arrows in *upper panel*). *Middle panel*, representative results of RT-PCR of RNA extracted from cells treated as follows: NT, not transfected; Ctrl, control siRNA as in microarray experiments; luc, luciferase; C1, C2, C3, *CELF1* siRNA 1 to 3; C123, mixture of 3 *CELF1* siRNAs; E1, E2, *ELAVL1* siRNA 1 and 2; E12, mixture of 2 *ELAVL1* siRNAs. *Lower panels*, we calculated the percentages of the isoform containing the exon of interest B. These percentages were divided by the corresponding ones from the untreated conditions (fold change, or FC) and are expressed as log_2_ ratios. The results of 3 independent experiments are shown. **B**, Same as **A**, but for *PHACTR2*, which contains a cassette exon B repressed by CELF1 but insensitive to ELAVL1 in microarray experiments. **C**, Same as **A** for *CD44* pre-mRNA, which contains a series of 4 cassette exons (v7 to v10) that have negative splicing indices after depletion of CELF1 or ELAVL1. C+E, mixture of 3 siRNAs against *CELF1* and 2 siRNAs against *ELAVL1*, as used in the microarray experiments. RT-PCR were done with primers located in exons labelled v6 and 15. *Lower panel*, we calculated the cumulative percentages of all isoforms containing the exons of interest: for exon v7, isoforms 1, 3, and 6; for v8 and v9, isoforms 1 to 4; and for exon v10, isoforms 1, 2, and 5.

Next, we used *PHACTR2* to investigate the specificity of the effect of the two depletions. *PHACTR2* mRNA encodes a phosphatase. It contains a cassette exon which is differentially included when CELF1 is depleted, but not with ELAVL1 depletion (Table S2). As with *WNK1*, the control siRNA does not modify *PHACTR2* splicing pattern, and the individual anti-*CELF1* and anti-*ELAVL1* siRNAs have the same effect as their combination (Figure 2B). However, as expected from the microarray data, when CELF1 (but not ELAVL1) is depleted, the long isoform containing the cassette exon is increased at the expense of the short one.

As an example of a more complex splicing pattern, we assessed *CD44* splicing. CD44 is a hyaluronan-binding signal-transducing cell surface receptor. *CD44* exons v2 to v10 are a series of cassette exons, and their inclusion or skipping produces many *CD44* isoforms that have different effects on cancer progression (Prochazka *et al*, 2014). Microarray hybridization showed that depletingCELF1 and/or ELAVL1 reduces the inclusion of exons v7 to v10, but this does not appear as significant for v7 and v9 after *p*-value adjustment (Table S2). The flanking exons v6 and 15 are similarly included in all conditions. Of the 16 isoforms which can theoretically be produced by the combinatorial inclusion or skipping of four exons, we detected seven using RT-PCR with primers located in exons v6 and 15 (Figure 2C). As above, *CD44’s* splicing pattern in non-transfected cells is very similar to that in cells transfected with either the control siRNA used in microarray experiments or an anti-luciferase siRNA (lanes 1, 2, 6). Importantly, in accordance with microarray data, all four alternative exons are repressed by the depletion of CELF1 or ELAVL1 (lanes 3-4), and are more strongly repressed by the simultaneous depletion of both proteins (lane 5). Together, these RT-PCR data on skipped exons all corroborate the microarray results.

Finally, we carried out RT-PCR validations on splicing events annotated as “alternative last exons” (Figure S2). *MICAL3, SPTBN1*, and *PPP3CB* encode a monooxygenase, a non-erythrocytic beta-II spectrin, and a calcium-dependent phosphatase, respectively. Each of these pre-mRNAs contains a terminal exon that can be also behave as an internal one, as it contains a 5’ splice site. Using this as a terminal exon results in a shorter C-terminal region in the encoded protein, with different subcellular localizations and molecular partners in the case of SPTBPN1 (Hayes *et al*, 2000). As expected from the microarray results (Table S2), the amounts of isoforms that result from using these as terminal exons are increased upon CELF1 depletion. They are also increased upon ELAVL1 depletion, but to a lesser extent (Figure S2). Altogether, these RT-PCR results (Figures 2 and S2) are fully in accordance with the microarray data (Table S2).

### CELF1 and ELAVL1 directly interact in the cell nucleus

Perhaps the joint regulation exerted by CELF1 and ELAVL1 on pre-mRNA processing is achieved by a macromolecular complex that contains these two proteins. The fact that they co-immunoprecipitate supports this interpretation (Le Tonquèze *et al*, 2010). We used a fluorescence lifetime imaging-based Förster Fluorescence Resonance Energy Transfer (FRET) approach to test whether the CELF1-ELAVL1 interaction is direct, and to identify the subcellular compartment in which it takes place. The donor lifetime decreases when FRET occurs, which reveals a close proximity between the donor (EGFP) and the acceptor (mCherry) (Padilla-Parra & Tramier, 2012). We expressed an EGFP-tagged version of CELF1 in HeLa cells. The nuclei of the cells expressing EGFP-CELF1 are green, with an EGFP fluorescence lifetime of about 2.5 ns (Figure 3). Those co-expressing an mCherry-tagged version of histone 2B (H2B) are yellow (Figure 3, merge channel), indicating a co-localisation of EGFP-CELF1 and mCherry-H2B in the nuclei of transfected cells. The unchanged lifetime of EGFP-CELF1 is consistent with a lack of interaction between CELF1 and H2B. In contrast, the lifetime of EGFP-CELF1 is reduced with a co-expressed mCherry-tagged version of CELF1, demonstrating the close proximity of EGFP and mCherry fluorophores, as expected due to CELF1’s previously demonstrated ability to oligomerise (Cosson *et al*, 2006). Importantly, mCherry-ELAVL1 also reduces the fluorescence lifetime of EGFP-CELF1 to a similar extent (Figure 3). Quantifying fluorescence lifetime in several nuclei demonstrated a highly significant decrease in the presence of mCherry-ELAVL1 (*p* < 2.2 × 10^−16^, Wilcoxon test) (Figure 3, lower panel). Swapping the two fluorophores produced the same results (Figure S3). We conclude therefore that CELF1 and ELAVL1 directly interact within a complex in the nucleus.

**Figure 3.**
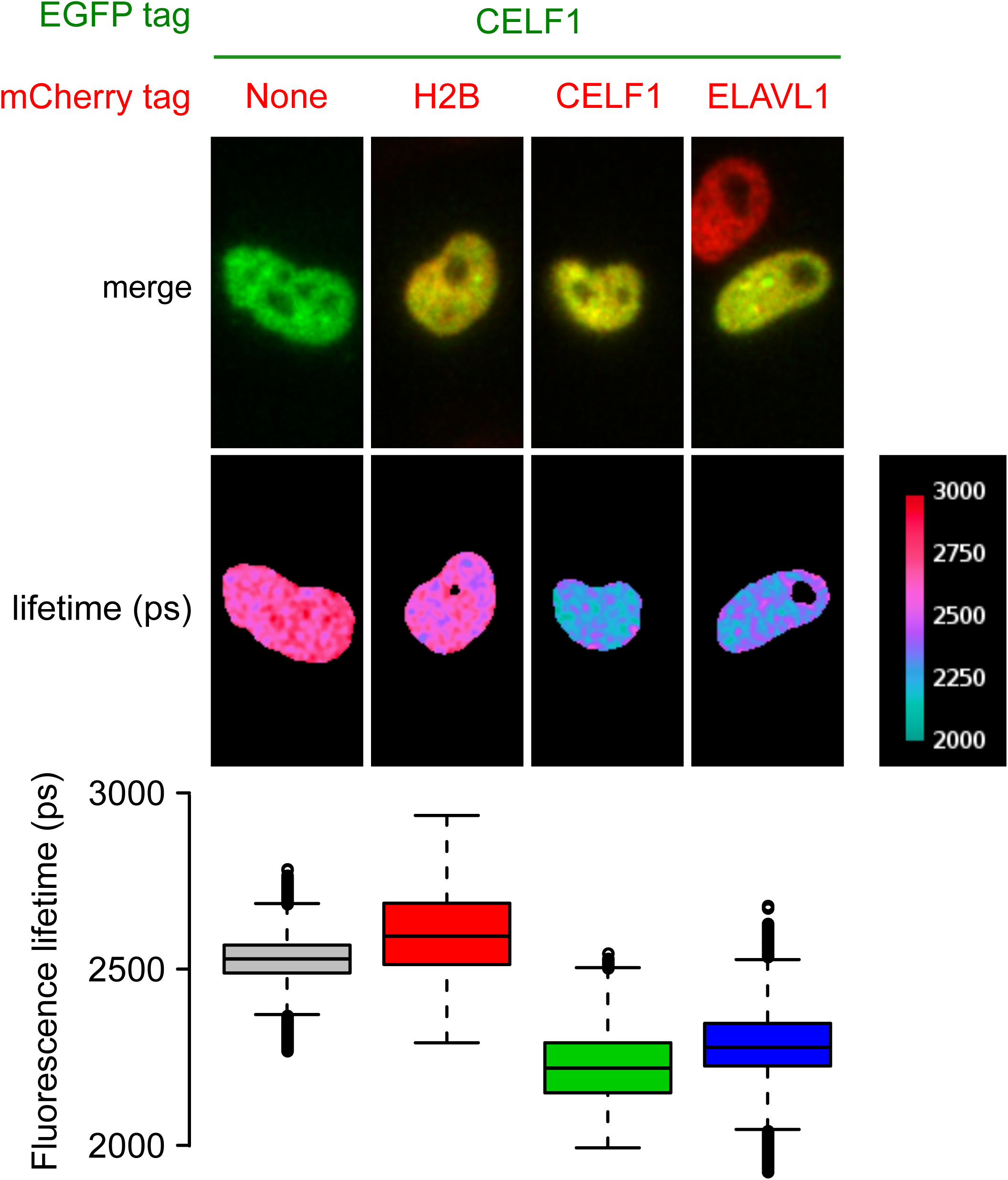
CELF1 and ELAVL1 directly interact in the nucleus. *Upper:* Representative images of HeLa cells co-transfected by a plasmid driving the expression of EGFP-tagged CELF1 and a plasmid that drives the expression of mCherry-tagged histone 2B (H2B), CELF1, or ELAVL1. The merge (EGFP plus mCherry) and EGFP lifetime channels are shown. The reduced EGFP lifetime in the two right panels implies that FRET occurs in these conditions. *Lower:* box plot of the distribution of EGFP fluorescence lifetimes in all measured pixels in 30 nuclei.

### *CELF1* and *ELAVL1* levels are correlated with human cancer prognosis

We show above that CELF1 and ELAVL1 control mRNA splicing together. Because HeLa cells come from cervical carcinoma, we used the TCGA (The Cancer Genome Atlas) uterine carcinosarcoma panel to test the relevance of the additive functions of these proteins in human pathology. Statistically, the survival rates of patients with either low and high *ELAVL1* mRNA levels are similar (Figure 4A), and the prognosis for patients with high *CELF1* levels is hardly better than that for those with low levels (*p* = 0.025, log-rank test, Figure 4B). Importantly, however, patients with low levels of both *CELF1* and *ELAVL1* have a worse survival rate (*p* = 9.7 × 10^−4^) than patients with a high level of *CELF1* and/or *ELAVL1* (Figure 4C). These data show that a more accurate prognosis can be given for uterine carcinosarcoma patients when both *CELF1* and *ELAVL1* mRNA levels are considered together rather than separately.

**Figure 4.**
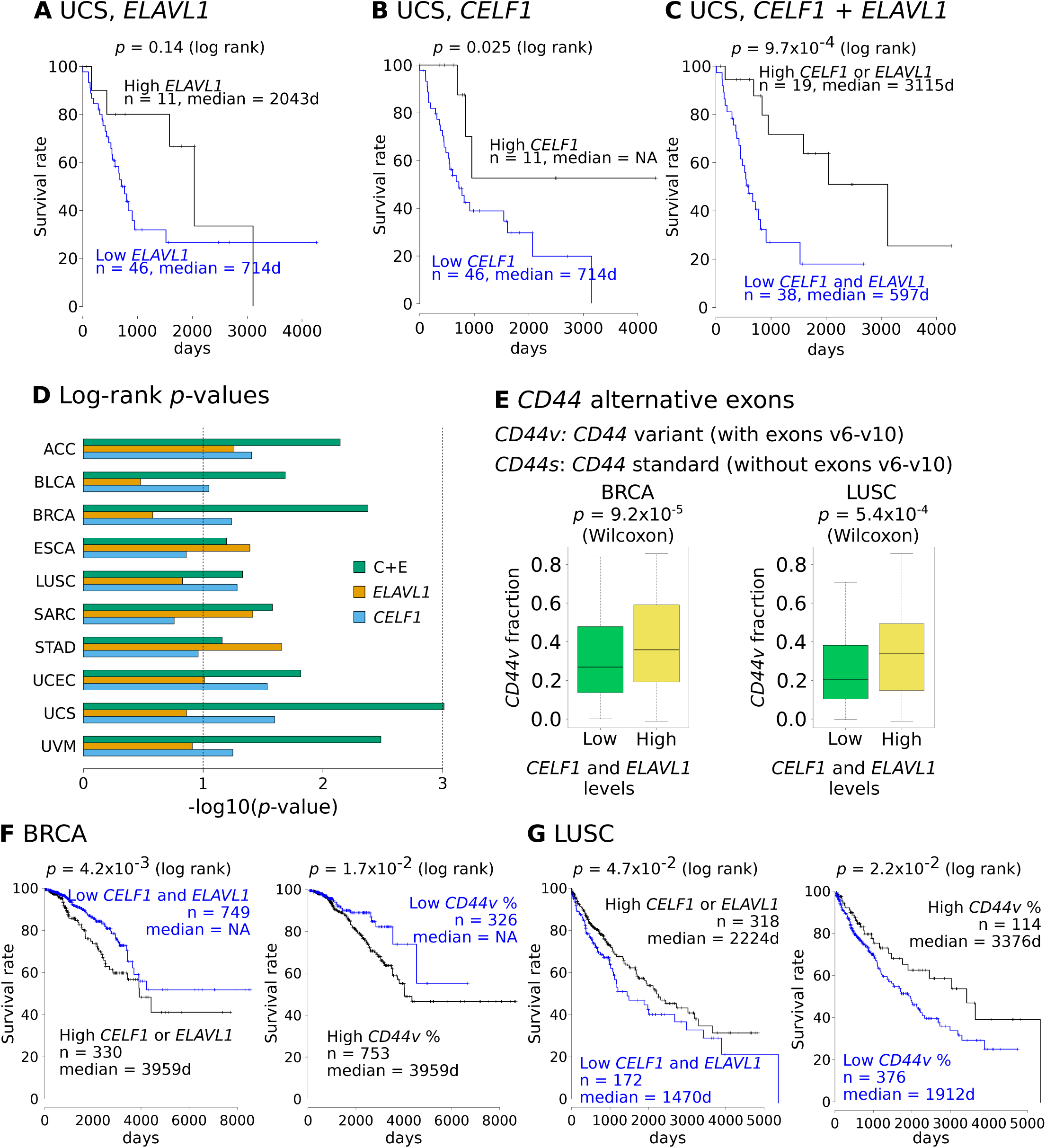
The combination of *CELF1* and *ELAVL1* is a prognostic marker in several cancers. **A**, Kaplan-Meier curve of low and high *ELAVL1* mRNA levels in 57 uterine carcinosarcoma patients (TCGA UCS panel). **B**, Same as **A** but for *CELF1* mRNA levels. **C**, Kaplan-Meier curve of the 57 patients showing low *CELF1* and *ELAVL1 m*RNA levels (n = 38, intersection of n = 46 low *CELF1* and n = 46 low *ELAVL1*), and high *CELF1* and/or *ELAVL1 m*RNA levels (n = 19, other patients). **D**, For the indicated TCGA cohorts, we calculated the log rank *p*-values for the difference in survival rates between low and high *CELF1* levels, low and high *ELAVL1* levels, and between low levels of both and high levels of at least one. The *p*-values are shown in a -log_10_ scale. ACC, adrenocortical carcinoma; BLCA, bladder urothelial carcinoma; BRCA, breast-invasive carcinoma; ESCA, esophageal carcinoma; LUSC, lung squamous cell carcinoma; SARC, sarcoma; STAD, stomach adenocarcinoma; UCEC, uterine corpus endometrial carcinoma; UCS, uterine carcinosarcoma; UVM, uveal melanoma. **E**, *Left*, Percentages of *CD44v* (including exons v6 to v10) in BRCA samples with expression levels of *CELF1* and *ELAVL1* in both the bottom 50% (n =254) and the top 50% (n = 253). *Right*, as left, but in LUSC samples, n = 125 for both groups. **F**, Kaplan-Meier curves of 1,079 BRCA patients according to either *CELF1* and *ELAVL1 m*RNA levels (*left*) or *CD44v* percentages (*right*). **G**, same as **F** but for 490 LUSC patients.

To expand these findings, we investigated the other TCGA cohorts. From them, we chose the cohorts where the effects of either low or high levels of *CELF1* and *ELAVL1* on patient prognosis are in the same direction, and removed those for which survival rates cannot be predicted (*p*> 0.1) by neither *CELF1* nor *ELAVL1* levels. We also removed those where the *CELF1* or *ELAVL1* levels alone are enough to separate the patients into two groups with different survival rates (*p* < 0.01 for at least one gene). For the ten resulting cohorts, we drew Kaplan-Meier curves for the levels of *CELF1, ELAVL1*, and both, and we compared the corresponding log-rank *p*-values (Figure 4D). For eight of the ten cohorts, we obtained lower *p*-values when looking at the levels of both *CELF1* and *ELAVL1*. The corresponding Kaplan-Meier curves are shown in Figure S4. These data indicate that the combined mRNA levels of *CELF1* and *ELAVL1* may be a valuable biomarker in a variety of human cancers.

*CD44* has a demonstrated role in cancer progression (Prochazka *et al*, 2014), and it is controlled by CELF1 and ELAVL1 at the splicing level. We therefore retrieved the data for the usage of *CD44* variable exons v6 to v10 (see Figure 2C) from TCGA SpliceSeq (Ryan *et al*, 2016). We investigated the eight above cohorts to see if there is a correlation between inclusion of these exons and the mRNA levels of *CELF1* and *ELAVL1*. Figure 4E shows that in breast invasive cancers (BRCA) and lung squamous cell carcinoma (LUSC), tumours having low levels of both *CELF1* and *ELAVL1* mRNA also have a significantly lower percentage of *CD44* isoforms containing exons v6 to v10 (*CD44v* or *CD44* variant). This is consistent with our results showing that CELF1 and ELAVL1 stimulate the inclusion of at least exons v7 to v10 in HeLa cells (Figure 2C). In breast invasive carcinoma, the survival rates are worse for patients with high *CELF1* and *ELAVL1* mRNA levels, as well as for those having a high percentage of *CD44v* (Figure 4F). Conversely, in lung squamous cell carcinoma, the survival rate is not worse but better when those same parameters are present (Figure 4G). Hence, although in both types of cancer the proteins CELF1 and ELAVL1 seem to promote *CD44v* by stimulating exon v7-v10 inclusion, the effects of this on patient prognosis are apparently completely different.

## Conclusion

Previous studies demonstrated an antagonism between ELAVL1 and CELF1 in translational control: in intestinal cells ELAVL1 stimulates the translation of *CDH1* (E-cadherin), *OCLN* (Occludin), and *MYC*, whereas CELF1 represses them (Yu *et al*, 2016; Liu *et al*, 2015; Yu *et al*, 2013). In contrast, here we see that these RNA-binding proteins actually cooperate, dictating mRNA splicing patterns. Our FRET experiments show that CELF1 and ELAVL1 are in direct contact in the nucleus, which supports the idea that they work together by belonging to the same macromolecular complex. This putative large complex may also include previously identified molecular partners of CELF1 such as HNRNPH1 and H2 (Paul *et al*, 2006), and/or partners of ELAVL1, such as HNRNPK (Hegele *et al*, 2012), HNRNPL (Matsui *et al*, 2008) and IGF2BP1 (Weidensdorfer *et al*, 2009).

Our findings considerably deepen the link between certain cancers and CELF1 and ELAVL1. It was already known that CELF1 protein levels are negatively correlated with patient survival in glioma and non-small-cell lung cancers (Xia *et al*, 2015; Jiao *et al*, 2013). This is also the case for ELAVL1 in ovarian high-grade serous carcinoma, colorectal cancer, and hepatocellular carcinoma (Davidson *et al*, 2016; Yoo *et al*, 2009; Zhu *et al*, 2015). We examined eight cohorts from a wide range of human tumours, and demonstrated that better prognostic values result from considering the expressions of both *CELF1* and *ELAVL1* than if these values are examined on their own. The cooperation between CELF1 and ELAVL1 in RNA splicing control provides a likely explanation for this.

One surprising observation is that high levels of *CELF1* and *ELAVL1* have opposite effects on patient survival in the presence of different types of cancer. BLCA, LUSC, UCS, or UVM patients who have high levels of *CELF1* and *ELAVL1* have better survival chances, but the opposite is true for patients with ACC, BRCA, SARC, or UCEC. An obvious explanation for this is that the transcriptomes of these tumours differ, as do the RNAs controlled by these two proteins. However, our *CD44* findings suggest an additional hypothesis. In breast cancer, a switch occurs during the epithelial-mesenchymal transition (EMT) from a *CD44v* isoform (containing exons v8-v10) to the standard *CD44s* isoform (devoid of the alternative cassette exons). This isoform switch is essential for EMT (Brown *et al*, 2011). *CD44* isoforms correlate with different cancer cell states, with *CD44s* associated with cell stemness, while *CD44v* is associated with cell proliferation (Zhang *et al*, 2019). Cell stemness and proliferation can both negatively impact patient survival, but perhaps in different ways for different tumour types. This would explain why *CD44v* percentages have contrary consequences for patient prognosis in breast and lung cancers (Figure 4). Globally, CELF1 and ELAVL1 may exert identical control of the same targets in different cancer types, but with disparate patient outcomes.

## MATERIALS AND METHODS

### Sequences and constructs

The sequences of the siRNAs and primers are given in the Supplemental material section. To construct plasmids EGFP-ELAVL1, mCherry-ELAVL1, EGFP-CELF1, and mCherry-CELF1, the cDNAs encoding ELAVL1 or CELF1 were introduced into the pENTR directional TOPO vector (Invitrogen), then inserted via Gateway recombination (Invitrogen) into the final 223 pCS EGFP DEST or 362 pCS Cherry DEST (Addgene) vectors.

### Cell manipulations and RNAi

We cultured HeLa Kyoto cells at 37 °C, 5% CO_2_ in DMEM (Gibco) with 10% fetal calf serum (PAA), 100 U/ml penicillin and 100 μg/ml streptomycin (Gibco). For RNAi, we used jetPRIME (Polyplus) to transfect the cells with the siRNAs indicated in the supplemental Material section. All experiments resulted in the same total concentration of 10 nM siRNAs. This was achieved by mixing the following siRNAs: 5 nM control plus 5 nM anti-ELAVL-1 siRNAs; 5 nM control plus 5 nM anti-CELF1 siRNAs; 5 nM anti-ELAVL-1 plus 5 nM anti-CELF1 siRNAs; 10 nM control siRNAs; and 10 nM anti-luciferase siRNAs. After 48 h, we recovered the cells for sampling in 500 μl of PBS by scraping after three washes with cold PBS then centrifuging for 5 min at 3000 rpm at 4 °C.

### FRET-FLIM

HeLa Kyoto cells were seeded at 20% confluence in observation chambers on 4-well glass slides (Dustcher). These were then transfected (jetPRIME) with 15 ng of the expression vectors of EGFP-labelled proteins in the presence or absence of 150 ng of vector that expresses mCherry-labelled proteins. After 24 h, the cells were washed with PBS and observed at 37 °C.

FRET experiments were conducted with a fastFLIM system using a plan APO 63x/1.4NA oil immersion objective (Leica). For EGFP donor excitation, a narrow 480/23 nm spectral band was selected from a Fianium white-light laser and sent to the CSUX1 microscope (Yokogawa) through a dichroic mirror (Di01-T405/488/568/647, Semrock) inside the spinning head. EGFP emissions were filtered on the spinning filter wheel (525/50 nm), then acquired with a time-gated intensified CCD camera (Picostar, LaVision or CoolSNAP HQ2, Roper). Five temporal gates of 2.25 ns were acquired sequentially by step-by-step adjustments of the laser signal delay to trigger the gated intensifier. Depending on the brightness of the sample, the CCD camera exposure time was between 500 ms and 2 s. The FLIM calculation to determine the mean fluorescence lifetime in a pixel-by-pixel basis was done online using flimager (Leray *et al*, 2013). For every nucleus region where lifetimes were measured, we checked for the absence of photo-bleaching and for the presence of both fluorophores for doubly transfected cells. Three independent series of transfection were carried out, with ten cells were analysed for each series, resulting in a total of 30 nuclei analysed per condition.

### Protein analysis

Cell pellets (equivalent to a 10-cm Petri dish with 80-90% confluence) were lysed with RIPA buffer (50 mM Tris-HCl pH 7.4, 150 mM NaCl, 1% sodium deoxycholate, 1% Triton X-100, 0.1% SDS, 0.1% P8340 Sigma protease inhibitor). After incubation for 10 min at room temperature, the samples were sonicated with a microprobe and centrifuged at 10,000 rpm for 5 min at 4 °C. Proteins were separated by electrophoresis and electrotransferred onto Hybond C Plus membranes (GE-Healthcare) per standard western blot procedures with anti-CELF1 (3B1 sc-20003, SantaCruz), anti-PCNA (p8825, Sigma-Aldrich), or anti ELAV1 (sc-5261, SantaCruz) primary antibodies, or with anti-mouse secondary antibodies. Imaging was done with a LI-COR Odyssey imager.

### RNA analysis

We treated the cells for 48 h with siRNAs. Three independent depletions were carried out for each condition. Total RNAs were extracted with TRIzol (Invitrogen) or a NucleoSpin RNA extraction kit (Macherey-Nagel), treated with Turbo DNase (Ambion), then sent to the Plateforme Génomique de Nantes for hybridization on SurePrint G3 human exon microarrays (Agilent). Raw signals were LOWESS-normalised before analysis. Alternatively, they were used as matrices to synthesize cDNA using random primers and SuperScript II reverse transcriptase (Invitrogen). PCR was done on cDNAs using unlabelled, Cy3-labelled, or ^32^P-labelled primers. Unlabelled RT-PCR (*PPP3CB, MICAL3*) results were analysed by SYBR staining after electrophoresis. The ^32^P-labelled RT-PCR (*CD44*) results were analysed by autoradiography. The Cy3-labelled RT-PCR (*WNK1, SPTBN1, PHACTR2*) results were treated as follows. DNA strands were prepared with a gene-specific forward primer having an M13 tail (CGCCAGGGTTTTCCCAGTCACGAC) for five cycles. These DNA strands were then used as PCR matrices with a Cy3-labelled M13 primer oligonucleotide and a gene-specific reverse primer. After electrophoresis, amplimere ratios were measured from the fluorescence intensities on a Typhoon imager (Amersham). In all the cases, the number of PCR cycles was determined empirically for each primer pair in order to remain within the exponential phase of amplification.

### Data analysis

The raw microarray data are available in GEO as data set GSE118981. For TCGA, we retrieved the data from Firebrowse (http://firebrowse.org/) and TCGA SpliceSeq (Ryan *et al*, 2016). We analysed all data using in-house R scripts, and these are available in the Supplemental material section.

## Supporting information

R script

## ACKNOWLEDGEMENTS

The results shown here are based in part on data generated by the TCGA Research Network (http://cancergenome.nih.gov/). Microarray hybridizations were carried out on the Plateforme Génomique de Nantes.

This research was funded by a grant from the Ligue contre le Cancer (comités 35, 22, 29) to LP. GD was supported by a joint doctoral fellowship from the Ligue contre le Cancer and by the Région Bretagne (ARED).

## AUTHOR CONTRIBUTIONS

GD performed and analysed the microarray experiments. GD, SPP, and MT designed, performed, and analysed the FRET experiments. SD, AM, WT and DR performed the depletions and RT-PCR. YA conceived and designed experiments and analysed data. LP conceived and designed experiments, analysed data, and wrote the manuscript.

## CONFLICT OF INTEREST

None.

## REFERENCES

Boulanger G, Cibois M, Viet J, Fostier A, Deschamps S, Pastezeur S, Massart C, Gschloessl B, Gautier-Courteille C & Paillard L (2015) Hypogonadism Associated with Cyp19a1 (Aromatase) Posttranscriptional Upregulation in Celf1 Knockout Mice. Mol. Cell. Biol. 35: 3244–3253

Brown RL, Reinke LM, Damerow MS, Perez D, Chodosh LA, Yang J & Cheng C (2011) CD44 splice isoform switching in human and mouse epithelium is essential for epithelial-mesenchymal transition and breast cancer progression. J Clin Invest 121: 1064–1074

Chang S-H, Elemento O, Zhang J, Zhuang ZW, Simons M & Hla T (2014) ELAVL1 regulates alternative splicing of eIF4E transporter to promote postnatal angiogenesis. Proc. Natl. Acad. Sci. U.S.A. 111: 18309–18314

Chaudhury A, Cheema S, Fachini JM, Kongchan N, Lu G, Simon LM, Wang T, Mao S, Rosen DG, Ittmann MM, Hilsenbeck SG, Shaw CA & Neilson JR (2016) CELF1 is a central node in post-transcriptional regulatory programmes underlying EMT. Nat Commun 7: 13362

Chen C-YA, Xu N & Shyu A-B (2002) Highly selective actions of HuR in antagonizing AU-rich element-mediated mRNA destabilization. Mol. Cell. Biol. 22: 7268–7278

Chi MN, Auriol J, Jégou B, Kontoyiannis DL, Turner JMA, de Rooij DG & Morello D (2011) The RNA-binding protein ELAVL1/HuR is essential for mouse spermatogenesis, acting both at meiotic and postmeiotic stages. Mol. Biol. Cell 22: 2875–2885

Cibois M, Boulanger G, Audic Y, Paillard L & Gautier-Courteille C (2012) Inactivation of the Celf1 gene that encodes an RNA-binding protein delays the first wave of spermatogenesis in mice. PLoS ONE 7: e46337

Cibois M, Gautier-Courteille C, Vallée A & Paillard L (2010) A strategy to analyze the phenotypic consequences of inhibiting the association of an RNA-binding protein with a specific RNA. RNA 16: 10–15

Cosson B, Gautier-Courteille C, Maniey D, Aït-Ahmed O, Lesimple M, Osborne HB & Paillard L (2006) Oligomerization of EDEN-BP is required for specific mRNA deadenylation and binding. Biol. Cell 98: 653–665

Damianov A, Ying Y, Lin C-H, Lee J-A, Tran D, Vashisht AA, Bahrami-Samani E, Xing Y, Martin KC, Wohlschlegel JA & Black DL (2016) Rbfox Proteins Regulate Splicing as Part of a Large Multiprotein Complex LASR. Cell 165: 606–619

Dassi E (2017) Handshakes and Fights: The Regulatory Interplay of RNA-Binding Proteins. Front Mol Biosci 4: 67

Davidson B, Holth A, Hellesylt E, Hadar R, Katz B, Tropé CG & Reich R (2016) HUR mRNA expression in ovarian high-grade serous carcinoma effusions is associated with poor survival. Hum. Pathol. 48: 95–101

Fialcowitz-White EJ, Brewer BY, Ballin JD, Willis CD, Toth EA & Wilson GM (2007) Specific protein domains mediate cooperative assembly of HuR oligomers on AU-rich mRNA-destabilizing sequences. J. Biol. Chem. 282: 20948–20959

Fu X-D (2004) Towards a splicing code. Cell 119: 736–738

García-Domínguez DJ, Morello D, Cisneros E, Kontoyiannis DL & Frade JM (2011) Stabilization of Dll1 mRNA by Elavl1/HuR in neuroepithelial cells undergoing mitosis. Mol. Biol. Cell 22: 1227–1239

Hall MP, Nagel RJ, Fagg WS, Shiue L, Cline MS, Perriman RJ, Donohue JP & Ares M (2013) Quaking and PTB control overlapping splicing regulatory networks during muscle cell differentiation. RNA 19: 627–638

Hayes NV, Scott C, Heerkens E, Ohanian V, Maggs AM, Pinder JC, Kordeli E & Baines AJ (2000) Identification of a novel C-terminal variant of beta II spectrin: two isoforms of beta II spectrin have distinct intracellular locations and activities. J. Cell. Sci. 113 (Pt 11): 2023–2034

Hegele A, Kamburov A, Grossmann A, Sourlis C, Wowro S, Weimann M, Will CL, Pena V, Lührmann R & Stelzl U (2012) Dynamic protein-protein interaction wiring of the human spliceosome. Mol. Cell 45: 567–580

Jiao W, Zhao J, Wang M, Wang Y, Luo Y, Zhao Y, Tang D & Shen Y (2013) CUG-binding protein 1 (CUGBP1) expression and prognosis of non-small cell lung cancer. Clin Transl Oncol 15: 789–795

Kalsotra A, Xiao X, Ward AJ, Castle JC, Johnson JM, Burge CB & Cooper TA (2008) A postnatal switch of CELF and MBNL proteins reprograms alternative splicing in the developing heart. Proc. Natl. Acad. Sci. U.S.A. 105: 20333–20338

Katsanou V, Milatos S, Yiakouvaki A, Sgantzis N, Kotsoni A, Alexiou M, Harokopos V, Aidinis V, Hemberger M & Kontoyiannis DL (2009) The RNA-binding protein Elavl1/HuR is essential for placental branching morphogenesis and embryonic development. Mol. Cell. Biol. 29: 2762–2776

Katz Y, Wang ET, Airoldi EM & Burge CB (2010) Analysis and design of RNA sequencing experiments for identifying isoform regulation. Nat. Methods 7: 1009–1015

Lafarga V, Cuadrado A, Lopez de Silanes I, Bengoechea R, Fernandez-Capetillo O & Nebreda AR (2009) p38 Mitogen-activated protein kinase- and HuR-dependent stabilization of p21(Cip1) mRNA mediates the G(1)/S checkpoint. Mol. Cell. Biol. 29: 4341–4351

Le Tonquèze O, Gschloessl B, Legagneux V, Paillard L & Audic Y (2016) Identification of CELF1 RNA targets by CLIP-seq in human HeLa cells. Genom Data 8: 97–103

Le Tonquèze O, Gschloessl B, Namanda-Vanderbeken A, Legagneux V, Paillard L & Audic Y (2010) Chromosome wide analysis of CUGBP1 binding sites identifies the tetraspanin CD9 mRNA as a target for CUGBP1-mediated down-regulation. Biochem. Biophys. Res. Commun. 394: 884–889

Leray A, Padilla-Parra S, Roul J, Héliot L & Tramier M (2013) Spatio-Temporal Quantification of FRET in living cells by fast time-domain FLIM: a comparative study of non-fitting methods [corrected]. PLoS ONE 8: e69335

Liu L, Ouyang M, Rao JN, Zou T, Xiao L, Chung HK, Wu J, Donahue JM, Gorospe M & Wang J-Y (2015) Competition between RNA-binding proteins CELF1 and HuR modulates MYC translation and intestinal epithelium renewal. Mol. Biol. Cell 26: 1797–1810

Masuda A, Andersen HS, Doktor TK, Okamoto T, Ito M, Andresen BS & Ohno K (2012) CUGBP1 and MBNL1 preferentially bind to 3’ UTRs and facilitate mRNA decay. Sci Rep 2: 209

Matsui K, Nishizawa M, Ozaki T, Kimura T, Hashimoto I, Yamada M, Kaibori M, Kamiyama Y, Ito S & Okumura T (2008) Natural antisense transcript stabilizes inducible nitric oxide synthase messenger RNA in rat hepatocytes. Hepatology 47: 686–697

Mukherjee N, Corcoran DL, Nusbaum JD, Reid DW, Georgiev S, Hafner M, Ascano M, Tuschl T, Ohler U & Keene JD (2011) Integrative regulatory mapping indicates that the RNA-binding protein HuR couples pre-mRNA processing and mRNA stability. Mol. Cell 43: 327–339

Padilla-Parra S & Tramier M (2012) FRET microscopy in the living cell: different approaches, strengths and weaknesses. Bioessays 34: 369–376

Pan Q, Shai O, Lee LJ, Frey BJ & Blencowe BJ (2008) Deep surveying of alternative splicing complexity in the human transcriptome by high-throughput sequencing. Nat. Genet. 40: 1413–1415

Paul S, Dansithong W, Kim D, Rossi J, Webster NJG, Comai L & Reddy S (2006) Interaction of muscleblind, CUG-BP1 and hnRNP H proteins in DM1-associated aberrant IR splicing. EMBO J. 25: 4271–4283

Peng SS, Chen CY, Xu N & Shyu AB (1998) RNA stabilization by the AU-rich element binding protein, HuR, an ELAV protein. EMBO J. 17: 3461–3470

Philips AV, Timchenko LT & Cooper TA (1998) Disruption of splicing regulated by a CUG-binding protein in myotonic dystrophy. Science 280: 737–741

Prochazka L, Tesarik R & Turanek J (2014) Regulation of alternative splicing of CD44 in cancer. Cell. Signal. 26: 2234–2239

Raghavachari N, Barb J, Yang Y, Liu P, Woodhouse K, Levy D, O’Donnell CJ, Munson PJ & Kato GJ (2012) A systematic comparison and evaluation of high density exon arrays and RNA-seq technology used to unravel the peripheral blood transcriptome of sickle cell disease. BMC Med Genomics 5: 28

Rahman MA, Masuda A, Ohe K, Ito M, Hutchinson DO, Mayeda A, Engel AG & Ohno K (2013) HnRNP L and hnRNP LL antagonistically modulate PTB-mediated splicing suppression of CHRNA1 pre-mRNA. Sci Rep 3: 2931

Rodan AR & Jenny A (2017) WNK Kinases in Development and Disease. Curr. Top. Dev. Biol. 123: 1–47

Roy A, Al-Qusairi L, Donnelly BF, Ronzaud C, Marciszyn AL, Gong F, Chang YPC, Butterworth MB, Pastor-Soler NM, Hallows KR, Staub O & Subramanya AR (2015) Alternatively spliced proline-rich cassettes link WNK1 to aldosterone action. J. Clin. Invest. 125: 3433–3448

Ryan M, Wong WC, Brown R, Akbani R, Su X, Broom B, Melott J & Weinstein J (2016) TCGASpliceSeq a compendium of alternative mRNA splicing in cancer. Nucleic Acids Res. 44: D1018–1022

Tang Y, Wang H, Wei B, Guo Y, Gu L, Yang Z, Zhang Q, Wu Y, Yuan Q, Zhao G & Ji G (2015) CUG-BP1 regulates RyR1 ASI alternative splicing in skeletal muscle atrophy. Sci Rep 5: 16083

Uren PJ, Burns SC, Ruan J, Singh KK, Smith AD & Penalva LOF (2011) Genomic analyses of the RNA-binding protein Hu antigen R (HuR) identify a complex network of target genes and novel characteristics of its binding sites. J. Biol. Chem. 286: 37063–37066

Vidal-Petiot E, Cheval L, Faugeroux J, Malard T, Doucet A, Jeunemaitre X & Hadchouel J (2012) A new methodology for quantification of alternatively spliced exons reveals a highly tissue-specific expression pattern of WNK1 isoforms. PLoS ONE 7: e37751

Vlasova IA, Tahoe NM, Fan D, Larsson O, Rattenbacher B, Sternjohn JR, Vasdewani J, Karypis G, Reilly CS, Bitterman PB & Bohjanen PR (2008) Conserved GU-rich elements mediate mRNA decay by binding to CUG-binding protein 1. Mol. Cell 29: 263–270

Wang ET, Sandberg R, Luo S, Khrebtukova I, Zhang L, Mayr C, Kingsmore SF, Schroth GP & Burge CB (2008) Alternative isoform regulation in human tissue transcriptomes. Nature 456: 470–476

Wang Z & Burge CB (2008) Splicing regulation: from a parts list of regulatory elements to an integrated splicing code. RNA 14: 802–813

Weidensdorfer D, Stöhr N, Baude A, Lederer M, Köhn M, Schierhorn A, Buchmeier S, Wahle E & Hüttelmaier S (2009) Control of c-myc mRNA stability by IGF2BP1-associated cytoplasmic RNPs. RNA 15: 104–115

Xia L, Sun C, Li Q, Feng F, Qiao E, Jiang L, Wu B & Ge M (2015) CELF1 is Up-Regulated in Glioma and Promotes Glioma Cell Proliferation by Suppression of CDKN1B. Int. J. Biol. Sci. 11: 1314–1324

Yoo PS, Sullivan CAW, Kiang S, Gao W, Uchio EM, Chung GG & Cha CH (2009) Tissue microarray analysis of 560 patients with colorectal adenocarcinoma: high expression of HuR predicts poor survival. Ann. Surg. Oncol. 16: 200–207

Yu T-X, Gu B-L, Yan J-K, Zhu J, Yan W-H, Chen J, Qian L-X & Cai W (2016) CUGBP1 and HuR regulate E-cadherin translation by altering recruitment of E-cadherin mRNA to processing bodies and modulate epithelial barrier function. Am. J. Physiol., Cell Physiol. 310: C54–65

Yu T-X, Rao JN, Zou T, Liu L, Xiao L, Ouyang M, Cao S, Gorospe M & Wang J-Y (2013) Competitive binding of CUGBP1 and HuR to occludin mRNA controls its translation and modulates epithelial barrier function. Mol. Biol. Cell 24: 85–99

Zhang H, Brown RL, Wei Y, Zhao P, Liu S, Liu X, Deng Y, Hu X, Zhang J, Gao XD, Kang Y, Mercurio AM, Goel HL & Cheng C (2019) CD44 splice isoform switching determines breast cancer stem cell state. Genes Dev. 33: 166–179

Zhang W, Yu Y, Hertwig F, Thierry-Mieg J, Zhang W, Thierry-Mieg D, Wang J, Furlanello C, Devanarayan V, Cheng J, Deng Y, Hero B, Hong H, Jia M, Li L, Lin SM, Nikolsky Y, Oberthuer A, Qing T, Su Z, et al (2015) Comparison of RNA-seq and microarray-based models for clinical endpoint prediction. Genome Biol. 16: 133

Zhu H, Berkova Z, Mathur R, Sehgal L, Khashab T, Tao R-H, Ao X, Feng L, Sabichi AL, Blechacz B, Rashid A & Samaniego F (2015) HuR Suppresses Fas Expression and Correlates with Patient Outcome in Liver Cancer. Mol. Cancer Res. 13: 809–818

