## Supplementary figures and images for "The RNA-binding proteins CELF1 and ELAVL1 cooperatively control RNA isoform production"

### Figure S1

David et al., Figure S1

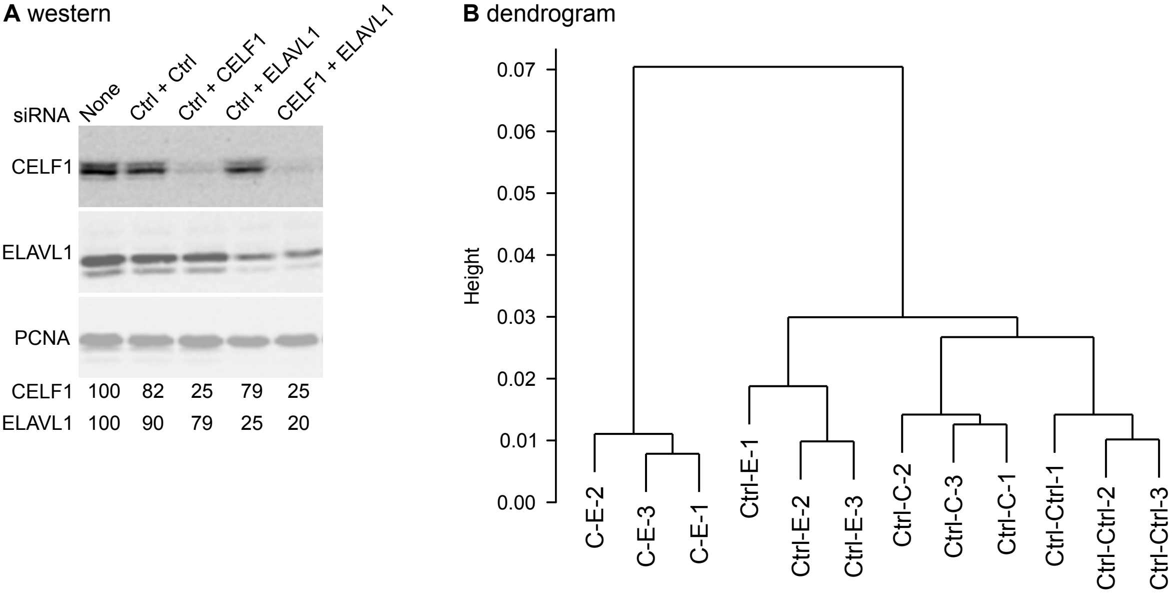

### Figure S2

David et al., Figure S2

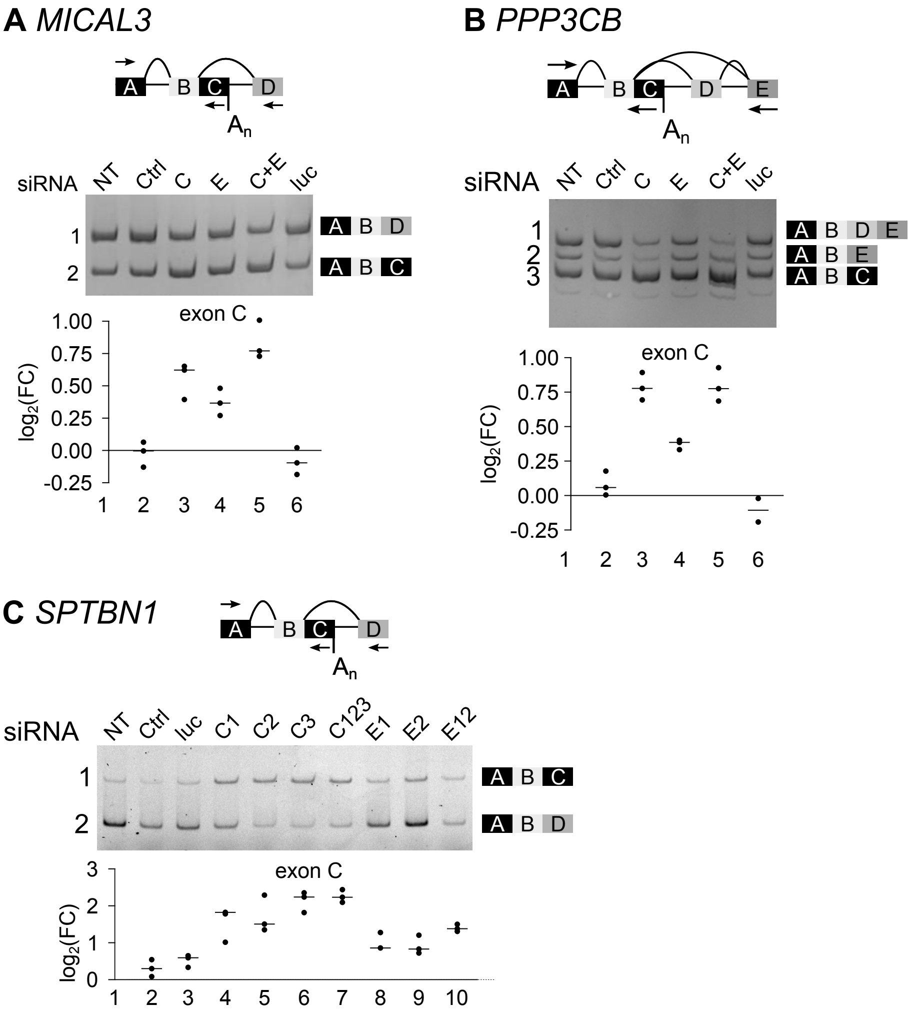

### Figure S3

David et al., Figure S3

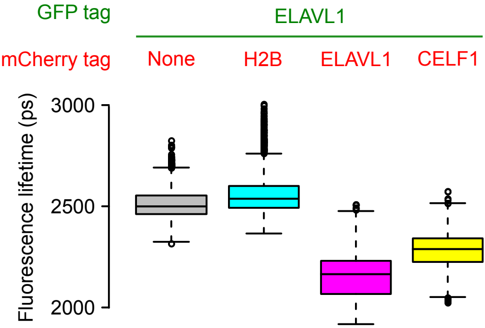

### Figure S4

#### David et al. Figure S4

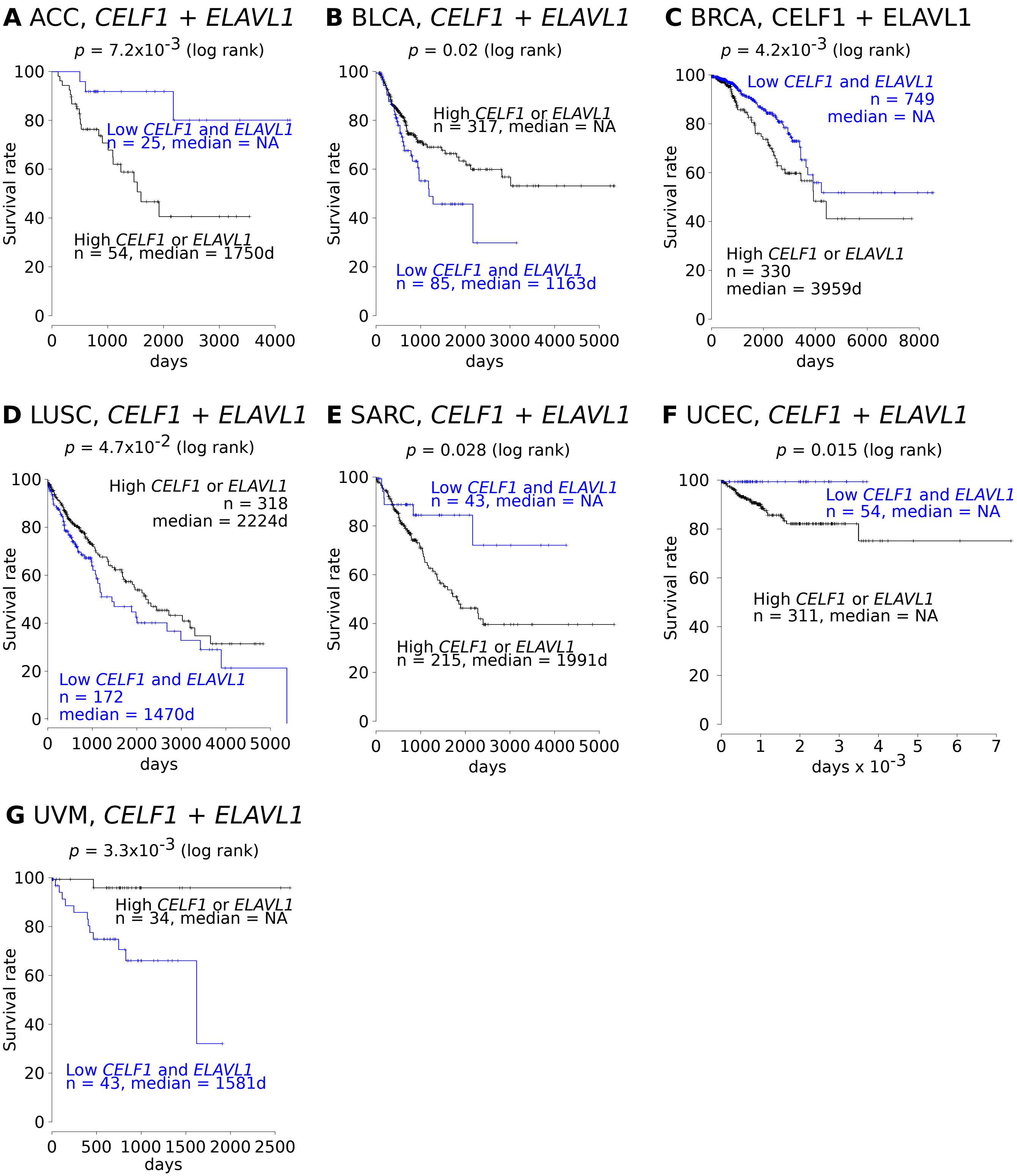
