## Supplemental text for "The RNA-binding proteins CELF1 and ELAVL1 cooperatively control RNA isoform production"

### SUPPLEMENTAL MATERIAL

#### SUPPLEMENTAL TABLE LEGENDS

**Table S1:** CELF1 and ELAVL1 ligands: list of genes (GeneName) that produce transcripts which interact with CELF1 and/or ELAVL1 in HeLa cells in CLIP-seq experiments (Le Tonquèze *et al*, 2016; Uren *et al*, 2011). n = 2,951 CELF1 ligands; n = 4,284 ELAVL1 ligands; and n = 1,475 ligands of both proteins.

**Table S2**: A table containing 62 rows and 20 columns. The rows represent the 62 exon array probes that in at least one test had significantly different normalized values (*FDR* = 0.1) than the values for the control condition (control siRNA-transfected cells). The columns are: GeneName; ProbeName; the normalized exon values for cells treated with 4 different siRNA combinations (control + control, control + CELF1, control + ELAVL1, and CELF1 + ELAVL1), each in triplicate; and finally the splicing indices and non-adjusted *p*-values for cells treated with control + CELF1, control + ELAVL1, and CELF1 + ELAVL1 siRNAs (each compared to control + control).

#### SUPPLEMENTAL FIGURE LEGENDS

**Figure S1. Quality control of microarray experiments**

**A**, Representative western blot showing the abundance of CELF1, ELAVL1, and PCNA (loading control) proteins in HeLa cells transfected with the indicated siRNA. The numbers beneath the gels indicate the protein amounts after normalisation by PCNA and the untransfected condition. **B,** Dendogram showing the hierarchical clustering of exon array probe values. We hybridised with exon arrays RNA extracted from cells transfected with the indicated combinations of siRNAs (C-E, *CELF1* and *ELAVL1* siRNAs; Ctrl-E, control and *ELAVL1* siRNAs; Ctrl-C, control and *CELF1* siRNAs; Ctrl-Ctrl, control siRNA). After normalisation and filtration for low signals, we retained 117,841 probes that correspond to exonic regions. Numbers 1-3 indicate independent siRNA transfections. The fact that identical combinations of siRNAs cluster together demonstrates that the results are reproducible.

**Figure S2. Examples of alternative last exons controlled by CELF1 and ELAVL1**

Representative RT-PCR experiments to measure the amounts of the isoforms of the *MICAL3* (**A**), *PPP3CB* (**B**), and *SPTBN1* (**C**) mRNAs, as per Figure 2. The *upper panels* show the pre-mRNA structure. Please note that for all three pre-mRNAs the "BC" exon can be used either as a terminal one, thanks to a cleavage/polyadenylation site (An), or as an internal one thanks to an internal 5' splice site between the B and C exonic regions. The isoform amounts were measured by RT-PCR, with one forward primer in exon A, one reverse primer in exonic region C (only amplifying the short isoform with the terminal BC exon), and one reverse primer in a downstream exon D or E (only amplifying the long isoform with the exons downstream of exon BC). The *middle panels* show a representative experiment. The *lower panels* show the quantification of three independent experiments. The 'C' exonic region has a significantly positive splicing index upon either CELF1 or ELAVL1 depletion in all three mRNAs, except in the case of ELAVL1 depletion in *PPP3CB*.

**Figure S3. EGFP-ELAVL1 fluorescence lifetime.**

We co-transfected HeLa cells with a plasmid driving the expression of EGFP-tagged ELAVL1 and plasmids driving the expression of mCherry-tagged histone 2B (H2B), CELF1, or ELAVL1. The quantifications of all measured pixels from 30 nuclei (three independent transfections) are shown as a box plot. A reduced EGFP fluorescence lifetime reveals that FRET occurs with mCherry, thus that the two fluorophores are in close proximity.

**Figure S4. Kaplan-Meier survival curves**

We show the Kaplan-Meier survival curves of patients grouped according to low *CELF1* and *ELAVL1* mRNA levels versus high *CELF1* or *ELAVL1* mRNA levels. The following cancers are shown: **A**, ACC, adrenocortical carcinoma; **B**, BLCA, bladder urothelial carcinoma; **C**, BRCA, breast invasive carcinoma; **D**, LUSC, lung squamous cell carcinoma; **E**, SARC, sarcoma; **F**, UCEC, uterine corpus endometrial carcinoma; **G**, UVM, uveal melanoma. The BRCA and LUSC data are the same as shown in Figures 4F-G.

#### SUPPLEMENTAL MATERIAL AND METHODS: SEQUENCES

##### Primer sequences

*WNK1* CTCCTCAACAGACAGTGCAG, GAAAGTACCCAGGTGTAGCCA

*PHACTR2* GAAAATTCAAACGGGCACAT, CCGGTGTTTCAGAGTGGTTT, CTTTGAAGCTTTGGGACGAG

*CD44* CAGAAGGAACAGTGGTTTGG, GGGTGGAATGTGTCTTGGTC

*MICAL3* TCTACTGTAAGCCACACTACTG, CAGCCAAACAAGGAAGTGAG, GTAGTTCTCCAGCTCGATCC

*PPP3CB* AAATTCGAGCAATTGGCAAG, AGACACTAGCTAATACAGTACCCTGTC, TGTGAGAGTCCCTGGGAAGT

*SPTBN1* AGCACGAAGGTTTCAGAGGA, GAACTTCACAGTCCAGACCA and TCTTGGCTTTACGATCAGAGG

**siRNA sequences**

*CELF1*: #1 GAGCCAACCUGUUCAUCUA, #2 GCUCUUUAUUGGUAUGAUU, #3 GCUGCAUUAGAAGCUCAGA

*ELAVL1*: #1 GAGGCAAUUACCAGUUUCA, #2 UCUUAAGUUUCGUAAGUUA

Ctrl: GUCUAGACGAGUGUGACAU

Luc: CAUUCUAUCCUCUAGAGGAUG
