## Supplementary material for "The RNA-binding proteins CELF1 and ELAVL1 cooperatively control RNA isoform production": R script

#################################################################################
###################### Prepare  files
#################################################################################
#ligands_celf.csv contains all the genes (genenames) with one CELF1 interacting cluster
#ligands_elav.csv contains all the genes (genenames) with one ELAVL1 interacting cluster
setwd("~/Luc_travail/Papiers/CEFL1-ELAVL1/manuscrit_aut18/R_script_and_results")
ligands_celf = read.table("ligands_celf.csv") #2951
ligands_elav = read.table("ligands_elav.csv") #4284
ligands = unique(rbind(ligands_celf, ligands_elav)) ##5760
# knowing that length(intersect(ligands_celf$V1, ligands_elav$V1)) == 1475 (ligands of both proteins) 
# 2951+4284-1475 == 5760, rbind did the job
ligands$celf_ligand = 0
ligands$elav_ligand = 0
ligands$both = 0
colnames(ligands)[1] = "GeneName"
head(ligands)
ligands$celf_ligand[ligands$GeneName %in% ligands_celf$V1] = 1
sum(ligands$celf_ligand) # 2951. OK
ligands$elav_ligand[ligands$GeneName %in% ligands_elav$V1] = 1
sum(ligands$elav_ligand) # 4284. OK
ligands$both = ligands$celf_ligand + ligands$elav_ligand
sum(ligands$celf_ligand == 1) #2951 CELF ligands
sum(ligands$celf_ligand == 0) #2809 non-CELF ligands (should be ligands of ELAV)
sum(ligands$elav_ligand == 1) #4284 ELAV ligands
sum(ligands$elav_ligand == 0) #1476 non-ELAV ligands (should be ligands of CELF)
sum(ligands$both == 2) #1475 ligands of both
sum(ligands$both == 0) # 0 ligands of none. All right.
write.table(ligands, file = "ligands", sep = "\t")
#The table "ligands" contains the genename, 2 columns with 0 or 1 to state if the gene is a CELF1 or ELAVL1 ligand, and a column with the sum of the 2 previous columns
# hence in ligands$both a "2" value flags the ligands of both ELAVL1 and CELF1.

# raw microarray data available in GEO as GSE118981
# The raw data were Lowess normalized => 233,164 probes
# Then filtration #1. For each probe threshold=average of medians of 12 samples. Keep probes that are above threshold at least twice about 3 in at least 1 condition
# Filtration #1 => 219,466
# Then averaging (arithmetic mean) when probes are more than once on the microarray => 122,123 probe values
# Then Filtration #2: remove probes not annotated as genic =>117,841 probes, in 17,806 genes

# Then prepare file data_exonnorm_SI_pval, including the splicing indexes and the pvalues for all the 117,841 probes
# This file is available in GEO as GSE118981_Processed_data.csv.gz
# start from valeurs_exons_normalisees.csv: normalized exon values for 117,841 probes.
# 14 columns=genename+probename+4 conditions (siScr, siCelf, siElav, siC+H), each in triplicate6 probes
setwd("~/Luc_travail/Papiers/CEFL1-ELAVL1/manuscrit_aut18/R_script_and_results")
data=read.table("valeurs_exons_normalisees.csv", h=T)
head(data)
Student = function(x)
{result=t.test(x[1:3], x[4:6])
 pvalue=result$p.value
 return(pvalue)
} #end of function
SI = function(x)
{result=mean(x[1:3])-mean(x[4:6])
 return(result)
} #end of function

#calculate the SI and pvalues for depletion of CELF1 vs mock (scr) depletion
data_celf_scr=as.matrix(data[ ,c(3:5, 12:14)]) #to make one dataframe with only the exon values for CELF1 and scr
pcelf=apply(data_celf_scr, 1, Student) 
length(pcelf) #117841 OK.
SI_celf=apply(data_celf_scr, 1, SI)
length(SI_celf) #117841 OK
# the same for ELAVL1 vs scr
data_elav_scr=as.matrix(data[ ,c(6:8, 12:14)]) #to make one dataframe with only the exon values for ELAVL1 and scr
pelav=apply(data_elav_scr, 1, Student) 
SI_elav=apply(data_elav_scr, 1, SI)
# the same for C+H vs scr
data_CH_scr=as.matrix(data[ ,c(9:11, 12:14)]) 
pCH=apply(data_CH_scr, 1, Student) 
SI_CH=apply(data_CH_scr, 1, SI)
# Make a new table
data_ext=cbind(data, SI_celf, pcelf, SI_elav, pelav, SI_CH, pCH)
head(data_ext)
write.table(data_ext, file = "data_exonnorm_SI_pval", sep = "\t")
#now we have a new file data_exonnorm_SI_pval.csv
# data_exonnorm_SI_pval.csv is a large file with 20 variables, 117841 observations (probes), in 17806 genes
# data_exonnorm_SI_pval.csv includes GeneName, ProbeName, Normalised exon values for SiScr, SiElav, SiCCelf, SiC+H, each in triplicate, splicing indexes and pvalues for  siCELF, siELav, siC+H

##I start from data_exonnorm_SI_pval.csv (same as GSE118981_Processed_data.csv.gz) to only retain the ligands
data = read.table("data_exonnorm_SI_pval", sep = "\t", h=T) #117841 probes in 17806 genes
ligands = read.table("ligands", sep = "\t")
ligands_both = ligands[ligands$both == 2, ]
data1 = data[data$GeneName %in% ligands_both$GeneName, ] #21146 probes
length(unique(data1$GeneName)) #1475 as expected for the number of ligands

#Then a filtration based on a reasonable level of expression of the genes (though the ligands are expected to be reasonably expressed)
# First I identify the expressed genes
# I start from "resultats_moy_geo_sur_sondes_filtrees.csv" = gene name+expressin in 17806 genes
toto = read.table("resultats_moy_geo_sur_sondes_filtrees.csv", h=T)
head(toto);dim(toto) #17806
# First the genes whose expression level is above first quartile in si_Celf in the three replicates
v1 = toto[,1] > quantile(toto[,1])[2] # a logical where TRUE is for "expressed" genes in IV_Celf
v2 = toto[,4] > quantile(toto[,4])[2] # a logical where TRUE is for "expressed" genes in III_Celf
v3 = toto[,8] > quantile(toto[,8])[2] # a logical where TRUE is for "expressed" genes in II_Celf
toto$expressed_in_sicelf = v1*v2*v3
sum(toto$expressed_in_sicelf) #12601 genes
# then the genes whose expression level is above first quartile in si_CH in the three replicates
v1 = toto[,2] > quantile(toto[,2])[2] # a logical where TRUE is for "expressed" genes in IV_CH
v2 = toto[,5] > quantile(toto[,5])[2] # a logical where TRUE is for "expressed" genes in III_CH
v3 = toto[,7] > quantile(toto[,7])[2] # a logical where TRUE is for "expressed" genes in II_Celf
toto$expressed_in_siCH = v1*v2*v3
sum(toto$expressed_in_siCH) #12790 genes
# then the genes whose expression level is above first quartile in si_SCR in the three replicates
v1 = toto[,3] > quantile(toto[,3])[2] # a logical where TRUE is for "expressed" genes in II_Scr
v2 = toto[,11] > quantile(toto[,11])[2] # a logical where TRUE is for "expressed" genes in III_Scr
v3 = toto[,12] > quantile(toto[,12])[2] # a logical where TRUE is for "expressed" genes in IV_Scr
toto$expressed_in_siScr = v1*v2*v3
sum(toto$expressed_in_siScr) #12605 genes
# then the genes whose expression level is above first quartile in si_HuR in the three replicates
v1 = toto[,6] > quantile(toto[,6])[2] # a logical where TRUE is for "expressed" genes in II_HuR
v2 = toto[,9] > quantile(toto[,9])[2] # a logical where TRUE is for "expressed" genes in III_HuR
v3 = toto[,10] > quantile(toto[,10])[2] # a logical where TRUE is for "expressed" genes in IV_HuR
toto$expressed_in_siHuR = v1*v2*v3
sum(toto$expressed_in_siHuR) #12573 genes
toto$expressed_somewhere = toto$expressed_in_sicelf + toto$expressed_in_siCH + toto$expressed_in_siScr + toto$expressed_in_siHuR
sum(toto$expressed_somewhere == 0) #4081 genes, which are considered as non expressed, 22.9%
sum(toto$expressed_somewhere > 0) #13725 "expressed" genes
expressed_genes = toto$GeneName[toto$expressed_somewhere > 0] #large factor 13725 elements

## now the filtration based on this criterion
# starting from data1, already filtered on the ligand criterion
data2 = data1[data1$GeneName %in% expressed_genes, ] #18727 probes
length(unique(data2$GeneName)) # 1299, to be compared with 1475, before filtration on expression level
#1299/1475 is 88% while 13725/17806 is only 77%. Hence while the expression-based filtration removes 23% of all genes, it only removes 12% of the ligands
# This was expected as ligands are supposed to be enriched in decently expressed genes
which(is.na(data2$GeneName)) #0
#remove the NA in the p values
length(which(is.na(data2$pCH))) # 13 rows to remove
data3 = data2[is.na(data2$pCH) == "FALSE", ] #18714 obs, which is 18727 - 13
write.table(data3, file = "probes_in_ligands", sep = "\t", row.names = F)

#multiple test adjustment (BH)
data3$pCH_BH = p.adjust (data3$pCH, method = "BH")
data3$pcelf_BH = p.adjust (data3$pcelf, method = "BH")
data3$pelav_BH = p.adjust (data3$pelav, method = "BH")
# I keep all the genes with FDR<0.1 in at least 1 condition
data4 = data3[data3$pCH_BH < 0.1 | data3$pcelf_BH < 0.1 | data3$pelav_BH < 0.1, ] #62 obs = 62 probes
length(unique(data4$GeneName)) # 57 = 57 genes, for 62 probes
write.table(data4[,1:20], file = "diff_probes_FDR<0.1", sep = "\t", row.names = F)


###################################################################################
################## Figure 1
###########################################################################
setwd("~/Luc_travail/Papiers/CEFL1-ELAVL1/manuscrit_aut18/R_script_and_results")


#############################################################
# MISO annotation
annotated_detected_events_117770=read.table("annotated_detected_events_117770")
head(annotated_detected_events_117770) #Column 2 is ProbeID (probe name) and Column 13 is the MISO splicing event associated with that probe (semicolon delimited is more than 1)
annotated_detected_events_117770=na.omit(annotated_detected_events_117770) #remove NA
typeof(annotated_detected_events_117770$detect_Alt_events) #integer. Why?? Transform to character
annotated_detected_events_117770$detect_Alt_events = as.character(annotated_detected_events_117770$detect_Alt_events)
#To get the number of probes per number of events
test=strsplit(annotated_detected_events_117770$detect_Alt_events, ";")
splicing_events=lapply(test, FUN=unique)
splicing_events_2=lapply(splicing_events, FUN=length)
summary(as.factor(unlist(splicing_events_2)))
# 0     1     2     3     4     5   6   7   8 (number of events per probe)
# 53036 36769 16743 6332  2427  796 132 22  9 (number of probes with this number of events)
#To get the numberof probes for each category of splicing event 
summary(as.factor(unlist(splicing_events)))
# A3SS      A5SS       AFE       ALE       MXE       RI        SE       TandemUTR 
# 15752     15846     13944      8070      3499      6170     38020      2656
#Now I start from diff_probes_FDR<0.1, which contains the 62 significant probes with FDR=0.1
data = read.table("diff_probes_FDR<0.1", sep = "\t", h=T) #62 obs of 20 variables
#I prepare 3 new matrices with the differential probes in SIcelf, SIelav, SICH
# SI-based filtration plus loose p-value filtration, because already done to prepare this data
diff_elav = data[abs(data$SI_elav)>0.2 & data$pelav < 0.05, ]  ##42 obs of 20 variables 
diff_celf = data[abs(data$SI_celf)>0.2 & data$pcelf < 0.05, ]  ##18 obs of 20 variables
diff_CH = data[abs(data$SI_CH)>0.2 & data$pCH < 0.05, ]  ## 60 obs of 20 variables 
names(splicing_events)=annotated_detected_events_117770$ProbeID
MISO_celf=splicing_events[names(splicing_events) %in% diff_celf$ProbeName] #list of 17
MISO_elav=splicing_events[names(splicing_events) %in% diff_elav$ProbeName] #list of 41
MISO_CH=splicing_events[names(splicing_events) %in% diff_CH$ProbeName] #list of 59
summary(as.factor(unlist(MISO_celf)))
#A3SS A5SS  AFE  ALE  MXE   SE 
#3    2    2    7    2   10 
summary(as.factor(unlist(MISO_elav)))
#A3SS      A5SS       AFE       ALE       MXE        SE TandemUTR 
#7         5         4        16         2        15         4 
summary(as.factor(unlist(MISO_CH)))
#A3SS      A5SS       AFE       ALE       MXE        SE TandemUTR 
#8         8         7        19         4        21         5
summary(as.factor(unlist(splicing_events)))
# A3SS      A5SS       AFE       ALE       MXE       RI        SE       TandemUTR 
# 15752     15846     13944      8070      3499      6170     38020      2656
#Now I have to merge all of this, but some columns are missing
table_MISO = matrix(nrow = 4, ncol = 8)
colnames(table_MISO) = c("A3SS", "A5SS", "AFE", "ALE", "MXE", "RI", "SE", "TandemUTR")
rownames(table_MISO) = c("all", "celf", "elav", "CH")
table_MISO[1, ] = unname(summary(as.factor(unlist(splicing_events))))
table_MISO[2, ] = c(unname(summary(as.factor(unlist(MISO_celf))))[1:5], 0, unname(summary(as.factor(unlist(MISO_celf))))[6], 0)
table_MISO[3, ] = c(unname(summary(as.factor(unlist(MISO_elav))))[1:5], 0, unname(summary(as.factor(unlist(MISO_elav))))[6:7])
table_MISO[4, ] = c(unname(summary(as.factor(unlist(MISO_CH))))[1:5], 0, unname(summary(as.factor(unlist(MISO_CH))))[6:7])
table_MISO
#     A3SS  A5SS   AFE  ALE  MXE   RI    SE TandemUTR
#all  15752 15846 13944 8070 3499 6170 38020      2656
#celf     3     2     2    7    2    0    10         0
#elav     7     5     4   16    2    0    15         4
#CH       8     8     7   19    4    0    21         5
# to prepare the barplot I put rows 2-4 in a single vector
toto = vector()
for (i in 1:8) {
  toto = c(toto, table_MISO[2,i], table_MISO[3,i], table_MISO[4,i])
} 
barplot(toto, horiz = T,  width = 1, xlim = c(0, 25), col = c(rgb(0, 0.6196, 0.451), rgb(0.8353, 0.3686, 0), rgb(0.9412, 0.8941, 0.2588)), space = 0) #colorblind friendly palette
#now calculate enrichment p values (hypergeometric test)
# first celf
table_MISO_pval = matrix(nrow = 3, ncol = 8)
for (i in 1:8)
{table_MISO_pval[1,i] = phyper(table_MISO[2,i], table_MISO[1,i], sum(table_MISO[1, ])-table_MISO[1,i], sum(table_MISO[2, ]), lower.tail = F)}
# then elav
for (i in 1:8)
{table_MISO_pval[2,i] = phyper(table_MISO[3,i], table_MISO[1,i], sum(table_MISO[1, ])-table_MISO[1,i], sum(table_MISO[3, ]), lower.tail = F)}
# then CH
for (i in 1:8)
{table_MISO_pval[3,i] = phyper(table_MISO[4,i], table_MISO[1,i], sum(table_MISO[1, ])-table_MISO[1,i], sum(table_MISO[4, ]), lower.tail = F)}
#then barplot
toto = vector()
for (i in 1:8) {
  toto = c(toto, table_MISO_pval[1,i], table_MISO_pval[2,i], table_MISO_pval[3,i])
} 
barplot(-log10(toto), horiz = T,  width = 1, xlim = c(0, 8), col = c(rgb(0, 0.6196, 0.451), rgb(0.8353, 0.3686, 0), rgb(0.9412, 0.8941, 0.2588)), space = 0) #colorblind friendly palette


##################################################
# SICelf vs SIelav for the differential probes (FDR=0.1)
data = read.table("diff_probes_FDR<0.1", sep = "\t", h=T) #62 obs of 20 variables
head(data)
# add MISO annotation
annotated_detected_events_117770=read.table("annotated_detected_events_117770")
head(annotated_detected_events_117770) #Column 2 is ProbeID (probe name) and Column 13 is the MISO splicing event associated with that probe (semicolon delimited is more than 1)
annotated_detected_events_117770=na.omit(annotated_detected_events_117770) #remove NA
annotated_detected_events_117770$detect_Alt_events = as.character(annotated_detected_events_117770$detect_Alt_events)
simplified_detected_events = annotated_detected_events_117770[ , c(2, 13)]
head(simplified_detected_events) #only keep probeID and alt events
subsel_MISO = simplified_detected_events[simplified_detected_events$ProbeID %in% data$ProbeName, ]
#61 obs instead of 62 in data, I lost one
data[which(data$ProbeName %in% subsel_MISO$ProbeID == F), ] 
#GeneName SLC30A7, ProbeName  A_37_P127875. Probe not found in annotated_detected_events. It's a retained intron, with positive splicing indices
colnames(subsel_MISO)[1] = "ProbeName" #to have the same name for the probe ID column in both tables
data_merge = merge(data, subsel_MISO, by = "ProbeName")
head(data_merge) #OK
#a function to get the genename and the MISO annotation
GeneAndAltEvent = function(dataset)
  {
  dataset$paste = paste(dataset$GeneName, "(", dataset$detect_Alt_events, ")", sep = " ")
  toto = as.character()
  limit = nrow(dataset)
  for (i in 1:limit ) 
    {
    toto = paste(toto, dataset$paste[i], sep = ", ")  
    } 
  return = toto  
  }
#list of genes with SI>0 for both CELF and ELAV
print(GeneAndAltEvent(dataset = data_merge[data_merge$SI_celf>0 & data_merge$SI_elav>0, ]))
# CCND1 (  ), TMPO ( ALE;ALE ), NEDD4L (  ), PCBP2 ( TandemUTR;TandemUTR;SE;SE ), SPOCK1 ( TandemUTR;TandemUTR ), PAFAH1B1 (  ), DDX46 (  ), HNRNPU ( ALE;A5SS;A5SS;A5SS;A5SS ), CPS1 ( TandemUTR;TandemUTR ), SETDB1 ( ALE;ALE;ALE;ALE ), SPTBN1 ( ALE;ALE;A5SS;A5SS;SE;SE;SE ), MICAL3 ( ALE;ALE;ALE ), SETDB1 ( ALE;ALE;ALE;ALE ), ITGA1 (  ), NAMPT ( TandemUTR;TandemUTR;ALE ), FANCA ( SE ), PDE4D ( ALE;ALE ), ANKRD17 (  ), HNRNPA1 ( ALE;ALE;ALE;A5SS;A5SS;A3SS;A3SS;SE;SE;SE;SE;SE;SE ), ITPR1 (  ), PKP4 ( ALE;ALE;ALE ), SBNO1 ( ALE ), WNK1 ( SE ), RABEP1 (  ), ZNF207 ( SE ), KLC1 ( ALE;ALE;ALE;ALE;ALE;ALE;SE;SE;SE ), PAPOLA ( ALE;ALE;SE;SE ), ATP2B4 ( ALE;A5SS;A5SS;SE;SE ), LPGAT1 (  ), CCBL2 ( A5SS;A5SS;SE;SE ), DST ( A3SS;A3SS ), PHACTR2 ( SE ), KIF13A ( AFE;AFE ), CREB3L2 ( ALE;ALE ), PDE7A ( ALE ), CELF1 ( A5SS;A5SS;A3SS;A3SS;SE;SE ), PPFIA1 ( ALE;SE;SE;SE;SE ), TOPBP1 ( ALE ), FANCA ( A3SS;A3SS;SE;SE ), RBMS1 ( AFE;AFE;AFE;A3SS;A3SS;SE;SE ), SLC30A9 (  )"
#n = 41
nrow(data[data$SI_celf > 0 & data$SI_elav > 0, ]) #42
# the difference is SLC30A7, which was removed for defective annotaiton in MISO yet has SI>0 for both CELF and ELAV
#list of genes with SI<0 for both CELF and ELAV
print(GeneAndAltEvent(dataset = data_merge[data_merge$SI_celf<0 & data_merge$SI_elav<0, ]))
# COL4A6 (  ), AHRR ( AFE;AFE;AFE;AFE;AFE;AFE ), TIA1 ( TandemUTR;TandemUTR;ALE ), MND1 (  ), SEPT9 ( AFE;AFE;AFE;AFE;AFE ), ASPH ( SE ), ANXA2 ( A5SS;A5SS;MXE;MXE;MXE;MXE;SE;SE ), CD44 ( MXE;SE;SE;SE ), TOPBP1 ( AFE;A3SS;A3SS ), IP6K2 ( MXE;MXE;SE;SE;SE ), TIA1 ( A3SS;A3SS ), ACOT9 ( A5SS;A5SS ), CENPP (  ), ALDH1A3 (  ), CD44 ( AFE;A3SS;A3SS;SE;SE;SE;SE ), C19orf42 ( AFE;SE;SE )" 
#n=16
nrow(data[data$SI_celf < 0 & data$SI_elav < 0, ]) #16. OK
#list of genes with SI<0 for CELF and SI>0 for ELAV
print(GeneAndAltEvent(dataset = data_merge[data_merge$SI_celf<0 & data_merge$SI_elav>0, ]))
#PRKD3 ( AFE )
nrow(data[data$SI_celf < 0 & data$SI_elav > 0, ]) #1 OK
#list of genes with SI>0 for CELF and SI<0 for ELAV
print(GeneAndAltEvent(dataset = data_merge[data_merge$SI_celf>0 & data_merge$SI_elav<0, ]))
# SLC7A6 (  ), COL4A5 (  ), MFF ( MXE;MXE;SE;SE;SE;SE;SE )"
nrow(data[data$SI_celf > 0 & data$SI_elav < 0, ]) #3. OK
# So I have 4 lists of genes, use Inkscape for a nicer display
#chi square test
toto = matrix(data = c(42, 3, 1, 16), nrow = 2, ncol = 2, byrow = T)
chisq.test(toto) #p-value = 2.092e-10


##################################################
## SIcelf + SIelav vs SI_CH for probes with FRD=0.1
data = read.table("diff_probes_FDR<0.1", sep = "\t", h=T) #62 obs of 20 variables
sum_SIC_SIH = data$SI_celf + data$SI_elav
data = cbind(data, sum_SIC_SIH)
head(data) ##OK
plot(data$sum_SIC_SIH ~ data$SI_CH, pch = 20, col = "grey"); abline(h=0);abline(v=0) 
correl=lm(data$sum_SIC_SIH ~ data$SI_CH) 
abline(correl) #regression line
summary(correl) ## R2 = 0.9309, p<2.2e-16
#Coefficients:
#Estimate Std. Error t value Pr(>|t|)    
#(Intercept)     0.09755    0.03264   2.988  0.00406 ** 
#data_sum$SI_CH   0.94869    0.03337  28.433  < 2e-16 ***
confint(correl, level = 0.99) #0.85993070 - 1.0374585. Confidence interval for the slope of the regression line
confint(correl, level = 0.95) # 0.88195199 1.015437. Confidence interval for the slope of the regression line
#to add MISO annotation
annotated_detected_events_117770=read.table("annotated_detected_events_117770")
annotated_detected_events_117770=na.omit(annotated_detected_events_117770) #remove NA
annotated_detected_events_117770$detect_Alt_events = as.character(annotated_detected_events_117770$detect_Alt_events)
simplified_detected_events = annotated_detected_events_117770[ , c(2, 13)]
head(simplified_detected_events) #only keep probeID and alt events
subsel_MISO = simplified_detected_events[simplified_detected_events$ProbeID %in% data$ProbeName, ]
subsel_ALE = data[data$ProbeName %in% subsel_MISO[grep("ALE", subsel_MISO$detect_Alt_events), 1],]
points(subsel_ALE$SI_CH, subsel_ALE$sum_SIC_SIH, col=rgb(0, 114, 178, maxColorValue = 255), pch=20) #to add ALE in blue = colorblinf friendly palette
subsel_SE = data[data$ProbeName %in% subsel_MISO[grep("SE", subsel_MISO$detect_Alt_events), 1],]
points(subsel_SE$SI_CH, subsel_SE$sum_SIC_SIH, col=rgb(213, 94, 0, maxColorValue = 255), pch=20) #to add the SE in vermillion = colorblinf friendly palette


###################################################################################
################## Analysis of RT-PCR (validations)
###########################################################################
setwd("~/Luc_travail/Papiers/CEFL1-ELAVL1/manuscrit_aut18/R_script_and_results")
data = read.csv("gel_quantification.csv", sep = ";", h = T, dec = ",")
head(data)
#column 1 is Gene (factor), column 2 is experiment number (factor), columns 3 to 24 are band intensities (numeric)
# For all conditions I compute SI, with respect to the "NT" conditions
#I start with the genes where I have C1, C2, C3, E1, E2, etc...
data1 = data[data$Gene %in% c("PHACTR2", "WNK1", "SPTBN1"), ]
data1$SI_Sc = log2((data1$Sc_included / (data1$Sc_included + data1$Sc_excluded)) / (data1$NT_included / (data1$NT_included + data1$NT_excluded)))
data1$SI_Luc = log2((data1$Luc_included / (data1$Luc_included + data1$Luc_excluded)) / (data1$NT_included / (data1$NT_included + data1$NT_excluded)))
data1$SI_C1 = log2((data1$siC1_included / (data1$siC1_included + data1$siC1_excluded)) / (data1$NT_included / (data1$NT_included + data1$NT_excluded)))
data1$SI_C2 = log2((data1$siC2_included / (data1$siC2_included + data1$siC2_excluded)) / (data1$NT_included / (data1$NT_included + data1$NT_excluded)))
data1$SI_C3 = log2((data1$siC3_included / (data1$siC3_included + data1$siC3_excluded)) / (data1$NT_included / (data1$NT_included + data1$NT_excluded)))
data1$SI_C123 = log2((data1$siC123_included / (data1$siC123_included + data1$siC123_excluded)) / (data1$NT_included / (data1$NT_included + data1$NT_excluded)))
data1$SI_E1 = log2((data1$siE1_included / (data1$siE1_included + data1$siE1_excluded)) / (data1$NT_included / (data1$NT_included + data1$NT_excluded)))
data1$SI_E2 = log2((data1$siE2_included / (data1$siE2_included + data1$siE2_excluded)) / (data1$NT_included / (data1$NT_included + data1$NT_excluded)))
data1$SI_E12 = log2((data1$siE12_included / (data1$siE12_included + data1$siE12_excluded)) / (data1$NT_included / (data1$NT_included + data1$NT_excluded)))
data1$SI_CE = log2((data1$siCE_included / (data1$siCE_included + data1$siCE_excluded)) / (data1$NT_included / (data1$NT_included + data1$NT_excluded)))
#build a new table to prepare the stripchart
SI_values = c(data1$SI_Sc, data1$SI_Luc, data1$SI_C1, data1$SI_C2, data1$SI_C3, data1$SI_C123, data1$SI_E1, data1$SI_E2, data1$SI_E12, data1$SI_CE)
SI_conditions = c(rep("Sc", times = nrow(data1)), rep("Luc", times = nrow(data1)), rep("C1", times = nrow(data1)), rep("C2", times = nrow(data1)), rep("C3", times = nrow(data1)), rep("C123", times = nrow(data1)), rep("E1", times = nrow(data1)), rep("E2", times = nrow(data1)), rep("E12", times = nrow(data1)), rep("CE", times = nrow(data1)))  
SI_gene = rep(c(rep("PHACTR2", times = 3), rep("WNK1", times = 3), rep("SPTBN1", times=3)), times = 10)
SI = data.frame(SI_values, as.factor(SI_conditions), as.factor(SI_gene))
colnames(SI) = c("values", "conditions", "gene")
SI$conditions = factor(SI$conditions, levels = c("Sc", "Luc", "C1", "C2", "C3", "C123", "E1", "E2", "E12", "CE")) #to have the plot in the same order as the figure
# Now plot the splicing indexes
SI1 = SI[SI$gene == "WNK1", ]
stripchart(values ~ conditions, data = SI1, vertical = T, pch=19)
abline(h=0, lty=2)
med = tapply(SI1$values, SI1$conditions, median)
segments(x0=seq(0.8, 9.8, by=1), y0=med, x1=seq(1.2, 10.2, by=1))
#then
SI1 = SI[SI$gene == "PHACTR2", ]
#and do it againt
SI1 = SI[SI$gene == "SPTBN1", ]

# Now the genes where I do not have C1, C2, C3, E1, E2, etc...
data1 = subset(data, !(data$Gene %in% c("PHACTR2", "WNK1", "SPTBN1")))
head(data1)
data1$SI_Sc = log2((data1$Sc_included / (data1$Sc_included + data1$Sc_excluded)) / (data1$NT_included / (data1$NT_included + data1$NT_excluded)))
data1$SI_Luc = log2((data1$Luc_included / (data1$Luc_included + data1$Luc_excluded)) / (data1$NT_included / (data1$NT_included + data1$NT_excluded)))
data1$SI_C123 = log2((data1$siC123_included / (data1$siC123_included + data1$siC123_excluded)) / (data1$NT_included / (data1$NT_included + data1$NT_excluded)))
data1$SI_E12 = log2((data1$siE12_included / (data1$siE12_included + data1$siE12_excluded)) / (data1$NT_included / (data1$NT_included + data1$NT_excluded)))
data1$SI_CE = log2((data1$siCE_included / (data1$siCE_included + data1$siCE_excluded)) / (data1$NT_included / (data1$NT_included + data1$NT_excluded)))
#build a new table to prepare the stripchart
SI_values = c(data1$SI_Sc, data1$SI_Luc, data1$SI_C123, data1$SI_E12, data1$SI_CE)
SI_conditions = c(rep("Sc", times = nrow(data1)), rep("Luc", times = nrow(data1)), rep("C123", times = nrow(data1)), rep("E12", times = nrow(data1)), rep("CE", times = nrow(data1)))  
SI_gene = rep(c(rep("CD44_exonb", times = 3), rep("CD44_exonc", times = 3), rep("CD44_exone", times=3), rep("PPP3CB", times=3), rep("JDP2", times=3), rep("ACSL4", times=4), rep("MICAL3", times=3), rep("CD44_small", times=3), rep("CD44_middle", times=3), rep("CD44_large", times=3)), times = 5)
SI = data.frame(SI_values, as.factor(SI_conditions), as.factor(SI_gene))
colnames(SI) = c("values", "conditions", "gene")
SI$conditions = factor(SI$conditions, levels = c("Sc", "C123", "E12", "CE", "Luc")) #to have the plot in the same order as the figure
# Now plot the splicing indexes
plotSI = function(gene) {
    SI1 = SI[SI$gene == gene, ]
    stripchart(values ~ conditions, data = SI1, vertical = T, pch=19, main = gene)
    abline(h=0, lty=2)
    med = tapply(SI1$values, SI1$conditions, median)
    segments(x0=seq(0.8, 4.8, by=1), y0=med, x1=seq(1.2, 5.2, by=1))
}
plotSI("CD44_large")
plotSI("CD44_small")
plotSI("CD44_middle")
plotSI("CD44_exonb")
plotSI("CD44_exonc")
plotSI("CD44_exone")
plotSI("PPP3CB")
plotSI("JDP2")
plotSI("MICAL3")

# a function to compute the splicing indexes from the quantifications of band intensities
compute_SI = function(dataset = data, gene = "PHACTR2")
{
  data1 = dataset[dataset$Gene == gene , ]
  NE_sc = data1$Sc_included / (data1$Sc_included + data1$Sc_excluded)  #normalised exon value for scrambled (control siRNA)
  SI_NT = log2((data1$NT_included / (data1$NT_included + data1$NT_excluded)) / NE_sc)
  SI_Luc = log2((data1$Luc_included / (data1$Luc_included + data1$Luc_excluded)) / NE_sc) 
  SI_C1 = log2((data1$siC1_included / (data1$siC1_included + data1$siC1_excluded)) / NE_sc) 
  SI_C2 = log2((data1$siC2_included / (data1$siC2_included + data1$siC2_excluded)) / NE_sc) 
  SI_C3 = log2((data1$siC3_included / (data1$siC3_included + data1$siC3_excluded)) / NE_sc) 
  SI_C123 = log2((data1$siC123_included / (data1$siC123_included + data1$siC123_excluded)) / NE_sc) 
  SI_E1 = log2((data1$siE1_included / (data1$siE1_included + data1$siE1_excluded)) / NE_sc) 
  SI_E2 = log2((data1$siE2_included / (data1$siE2_included + data1$siE2_excluded)) / NE_sc) 
  SI_E12 = log2((data1$siE12_included / (data1$siE12_included + data1$siE12_excluded)) / NE_sc) 
  SI_CE = log2((data1$siCE_included / (data1$siCE_included + data1$siCE_excluded)) / NE_sc) 
  tmp = data.frame(gene, mean(SI_NT), mean(SI_Luc, na.rm = T), mean(SI_C1), mean(SI_C2), mean(SI_C3), mean(SI_C123), mean(SI_E1), mean(SI_E2), mean(SI_E12), mean(SI_CE), sd(SI_NT), sd(SI_Luc, na.rm = T), sd(SI_C1), sd(SI_C2), sd(SI_C3), sd(SI_C123), sd(SI_E1), sd(SI_E2), sd(SI_E12), sd(SI_CE))
  #na.rm = T because 1 miss 1 value for luc
  colnames(tmp) = c("Gene", "mean_NT", "mean_Luc", "mean_C1", "mean_C2", "mean_C3", "mean_C123", "mean_E1", "mean_E2", "mean_E12", "mean_CE", "sd_NT", "sd_Luc", "sd_C1", "sd_C2", "sd_C3", "sd_C123", "sd_E1", "sd_E2", "sd_E12", "sd_CE")
  return = tmp
}
alt_compute_SI = function(dataset = data, gene = "JDP2")
{
  data1 = dataset[dataset$Gene == gene , ]
  NE_sc = data1$Sc_included / data1$Sc_excluded  # for JDP2 excluded is in fact total
  SI_NT = log2((data1$NT_included / data1$NT_excluded) / NE_sc)
  SI_Luc = log2((data1$Luc_included / data1$Luc_excluded) / NE_sc) 
  SI_C1 = log2((data1$siC1_included / data1$siC1_excluded) / NE_sc) 
  SI_C2 = log2((data1$siC2_included / data1$siC2_excluded) / NE_sc) 
  SI_C3 = log2((data1$siC3_included / data1$siC3_excluded) / NE_sc) 
  SI_C123 = log2((data1$siC123_included / data1$siC123_excluded) / NE_sc) 
  SI_E1 = log2((data1$siE1_included / data1$siE1_excluded) / NE_sc) 
  SI_E2 = log2((data1$siE2_included / data1$siE2_excluded) / NE_sc) 
  SI_E12 = log2((data1$siE12_included / data1$siE12_excluded) / NE_sc) 
  SI_CE = log2((data1$siCE_included / data1$siCE_excluded) / NE_sc) 
  tmp = data.frame(gene, mean(SI_NT), mean(SI_Luc), mean(SI_C1), mean(SI_C2), mean(SI_C3), mean(SI_C123), mean(SI_E1), mean(SI_E2), mean(SI_E12), mean(SI_CE), sd(SI_NT), sd(SI_Luc), sd(SI_C1), sd(SI_C2), sd(SI_C3), sd(SI_C123), sd(SI_E1), sd(SI_E2), sd(SI_E12), sd(SI_CE))
  colnames(tmp) = c("Gene", "mean_NT", "mean_Luc", "mean_C1", "mean_C2", "mean_C3", "mean_C123", "mean_E1", "mean_E2", "mean_E12", "mean_CE", "sd_NT", "sd_Luc", "sd_C1", "sd_C2", "sd_C3", "sd_C123", "sd_E1", "sd_E2", "sd_E12", "sd_CE")
  return = tmp
}
# run compute_psi for all the genes in data
SI_PHACTR2 = compute_SI(dataset = data, gene = "PHACTR2")
SI_WNK1 = compute_SI(dataset = data, gene = "WNK1")
SI_SPTBN1 = compute_SI(dataset = data, gene = "SPTBN1")
SI_CD44_exonb = compute_SI(dataset = data, gene = "CD44_exonb")
#significant in microarray
SI_CD44_exonc = compute_SI(dataset = data, gene = "CD44_exonc") ##the same as exon d (sames bands include or exclude exons c and d.
# exon d is the significant one in microarray, replace exon c by exond 
SI_CD44_exone = compute_SI(dataset = data, gene = "CD44_exone")
#not significant in microarray
SI_PPP3CB = compute_SI(dataset = data, gene = "PPP3CB")
#caution, for JDP2 the other band is not excluded but total. I have to run alt_compute_SI
SI_JDP2 = alt_compute_SI(dataset = data, gene = "JDP2")
computed_SI = rbind(SI_PHACTR2, SI_WNK1, SI_SPTBN1, SI_CD44_exonb, SI_CD44_exonc, SI_CD44_exone, SI_PPP3CB, SI_JDP2)

# a function to barplot the SI, with SD
barplot_SI = function(dataset = computed_SI, gene = "PHACTR2", ylim = c(-2, +2))
{
  data1 = dataset[dataset$Gene == gene , ]
  meanval = as.matrix(data1[2:11])
  sdval = as.matrix(data1[12:21])
  bp = barplot(meanval, ylim = ylim, ylab = "splicing index", main = gene, col = "grey")
  arrows(bp, meanval-sdval, bp, meanval+sdval, lwd=1.5, angle=90, length=0.1, code=3) 
  abline(h = 0)
}
#plots
barplot_SI(dataset = computed_SI, gen = "PHACTR2", ylim = c(-2, 3))
barplot_SI(dataset = computed_SI, gen = "WNK1", ylim = c(-1.5, 3))
barplot_SI(dataset = computed_SI, gen = "SPTBN1", ylim = c(-1, 3))
barplot_SI(dataset = computed_SI, gen = "CD44_exonb", ylim = c(-3, 1))
barplot_SI(dataset = computed_SI, gen = "CD44_exonc", ylim = c(-3, 1))
barplot_SI(dataset = computed_SI, gen = "CD44_exone", ylim = c(-3, 1))
barplot_SI(dataset = computed_SI, gen = "PPP3CB", ylim = c(-0.6, 1))
barplot_SI(dataset = computed_SI, gen = "JDP2", ylim = c(-0., 2.5))


##############################################################
########  TCGA 
#####################################################################
### First I download the gene expression data in different cohorts from Firebrowse
## Look like "BRCA.rnaseqv2__illuminahiseq_rnaseqv2__unc_edu__Level_3__RSEM_genes_normalized__data.data.txt"

####### A function to get the RNA levels (normalised RSEM) in each sample for the genes in "gene_set"
gene_levels = function(data_genexp, gene_set)
{
  data = data_genexp
  gene_set = unique(as.character(gene_set))
  gene_set_1 = c(gene_set, "Hybridization REF") #this will permit to keep the patient IDs in the simplified table
  data[, 1] = as.character(data[,1])
  data[,1] = gsub("|", "asterix",  data[,1], fixed=T) #fixed = T no regexp
  data[,1] = gsub("asterix.{1,}", "", data[,1]) #gsub in 2 passes as I both need a regexp and | not to be seen as a regexp 
  data1 = data[data[,1] %in% gene_set_1, ]
  data2 = as.data.frame(t(data1))
  data3 = data.frame(unname(data2[2:nrow(data2), 1])) #patients ID
  for (i in 1:length(gene_set))
  {
    data_tmp = data.frame(as.numeric(as.character(unname(data2[2:nrow(data2), i+1])))) 
    data3 = data.frame(data3, data_tmp)
  }
  colnames(data3) = t(data2[1,])
  colnames(data3) = paste(colnames(data3), "_level", sep ="")
  colnames(data3)[1]= "IDs"
  return = data3
}

###### A function to add the type (tumor or normal) to the table of gene level
add_type = function(data_gene)
{
  type = vector(length = nrow(data_gene))  
  type[grep("-11A-", data_gene$IDs)] = "Norm"
  type[grep("-01A-", data_gene$IDs)] = "Tumor"
  type[grep("-01B-", data_gene$IDs)] = "Tumor"
  type[grep("-03A-", data_gene$IDs)] = "Tumor"
  type[grep("-03B-", data_gene$IDs)] = "Tumor"
  #norm samples are indexed by 11A in firebrowse, tumor samples are indexed by 01A
  data_gene_1 = cbind(data_gene, as.factor(type))
  colnames(data_gene_1)[ncol(data_gene_1)] = "type"
  data_gene_1 = data_gene_1[data_gene_1$type != "FALSE",]
  data_gene_1$type = droplevels(data_gene_1$type)
  return = data_gene_1
}

####### A function to have the splice data from TCGA splice seq for a limited number of genes
# Files downloaded from TCGA spliceseq contain all the splice events relative to a list of genes
PSI_goi = function(PSI_download, gene_set)
{
  PSI_download_1 = PSI_download[PSI_download$symbol %in% gene_set, ]
  return = PSI_download_1
}

##### A function to only retain splicing events of interest
##### Has to be done manually...
clean_PSI_selected = function(PSI_CELFELAV)
{
  PSI = PSI_CELFELAV
  PSI = PSI[PSI$symbol != "CCBL2" | PSI$exons == 2 , ] #CCBL2: exon 2 is ES. I keep the genes different from CCBL2 or whose "exon" is exon 2
  PSI = PSI[PSI$symbol != "SLC30A7", ]  #SLC30A7 not clear what happens in TCGA spliceseq. Remove the gene
  PSI = PSI[PSI$symbol != "SETDB1" | PSI$exons == 9.2 , ]  #SETDB1: exon9.2 is alternative terminal exon
  PSI = PSI[PSI$symbol != "ATP2B4", ]  #ATP2B4: event not in TCGA spliceseq
  PSI = PSI[PSI$symbol != "HNRNPU", ]  #HNRNPU: event not in TCGA spliceseq
  PSI = PSI[PSI$symbol != "CPS1", ]  #ceseq
  PSI = PSI[PSI$symbol != "CD44" | PSI$exons == "10:11:12.1:13:14" , ] #CD44: it's a mess.. I take globally the event "10:11:12.1:13:14", which considers the inclusion or exclusion of 5 consecutive exons that all are down in depletion
  PSI = PSI[PSI$symbol != "CELF1", ]  #CEFL1: I prefer remove it as it might interfere with the CELF1 level
  PSI = PSI[PSI$symbol != "PPFIA1" | PSI$exons == "27.2" , ]  #PPFIA1: exon of interest is 27.2 (distal region of a composite internal-terminal exon)
  PSI = PSI[PSI$symbol != "WNK1" | (PSI$exons == "14" & PSI$to_exon == "15"), ] #WNK1: exon of insterest is #14
  PSI = PSI[PSI$symbol != "PCBP2" | PSI$exons == "18" , ]#PCBP2: alternative termionation, exon 18
  PSI = PSI[PSI$symbol != "HNRNPA1", ]  #HNRNPA1: alternative termination, nothing in TCGA splice seq
  PSI = PSI[PSI$symbol != "TMPO" | PSI$exons == "4" , ] #TMPO: alternative terminal exon numbered 4
  PSI = PSI[PSI$symbol != "PAPOLA" | PSI$exons == "7.2" , ]  #PAPOLA: exon 7.2 is th edistal part of a composite exon (up in KD)
  PSI = PSI[PSI$symbol != "KLC1", ]   #KLC1: alternative terminal exon. TCGA splice seq annotation not consistent
  PSI = PSI[PSI$symbol != "ANXA2" | PSI$to_exon == "3" , ]   #ANXA2: stimulated ES #2 but "PSI$exons == "3"" retains 2 events 
  PSI = PSI[PSI$symbol != "SLC7A6" | PSI$exons == "12" , ] #SLC7A6: skipped exon #12
  PSI = PSI[PSI$symbol != "FANCA", ]  #FANCA : exon 4.2 annotated as AT whereas it's ES. Unclear
  PSI = PSI[PSI$symbol != "PAFAH1B1", ]  #PAFAH1B1: tandfem CPA non annotated in TCGA spliceqes
  PSI = PSI[PSI$symbol != "RABEP1" | PSI$exons == "4" , ]   #RABEP1: skipped exon #4
  PSI = PSI[PSI$symbol != "ZNF207" | PSI$exons == "10" , ] #ZNF207: skipped exon #10
  PSI = PSI[PSI$symbol != "NEDD4L", ] #NEDD4L tandem CPA not annotated in TCGA spliceqes
  PSI = PSI[PSI$symbol != "SPTBN1" | PSI$exons == "33.2" , ]  #SPTBN1: composite internal-terminal, distal part (used as terminal) is 33.2
  PSI = PSI[PSI$symbol != "TIA1", ]     #TIA1: event unclear
  PSI = PSI[PSI$symbol != "PKP4", ]   #PKP4 not consistent with TCGZA splice seq
  PSI = PSI[PSI$symbol != "RBMS1", ]   #RBMS1 SE, an exon between the 1st and the 3rd which is not annotated #2 in TCGA splcie seq
  PSI = PSI[PSI$symbol != "MICAL3" | PSI$exons == "25.2" , ]  #MICAL3 alternative last exon #25.2
  PSI = PSI[PSI$symbol != "ITPR1", ]  #ITPR1 it's a mess!!
  PSI = PSI[PSI$symbol != "IP6K2" | PSI$exons == "11.9" , ]  #IP6K2 alternative last exon 11.9
  PSI = PSI[PSI$symbol != "SLC30A9", ]  #SLC30A9 the differentiel exon is not annotated as alternative in TCGA spliceseq
  PSI = PSI[PSI$symbol != "ANKRD17", ]  #ANKRD17 the differentiel exon is not annotated as alternative in TCGA spliceseq
  PSI = PSI[PSI$symbol != "MND1" | PSI$exons == "1" , ]  #MND1 alternative promotor exon 1
  PSI = PSI[PSI$symbol != "AHRR", ]   #AHRR the differentiel exon is not annotated as alternative in TCGA spliceseq
  PSI = PSI[PSI$symbol != "PDE4D", ]   #PDE4D tandem CPA not annotated in TCGA spliceseq
  PSI = PSI[PSI$symbol != "DDX46", ]  #DDX46 tandem CPA not annotated in TCGA spliceseq
  PSI = PSI[PSI$symbol != "DST", ]   #DST: too many exons to understand anything
  PSI = PSI[PSI$symbol != "PHACTR2" | PSI$exons == "8" , ]  #PHACTR2: SE#8
  PSI = PSI[PSI$symbol != "NAMPT", ]   #NAMPT: alternative tandem CPA not annotated in TCGA spliceseq
  PSI = PSI[PSI$symbol != "LPGAT1", ]   #NAMPT: alternative tandem CPA not annotated in TCGA spliceseq
  PSI = PSI[PSI$symbol != "CREB3L2", ]   #CREB3L2 unclear
  PSI = PSI[PSI$symbol != "ASPH" | PSI$exons == "12" , ]   #ASPH: skipped exon #12 (SI<0)
  PSI = PSI[PSI$symbol != "PDE7A", ]  #PDE7A tandem not annotated in TCGA splice seq
  PSI = PSI[PSI$symbol != "ACOT9" , ]  #From microarray data, exons downstream of exon 14 are down, and in UCSC exon 14 can be the first promoter, but not in TCGA spliceqes 
  PSI = PSI[PSI$symbol != "COL4A6", ]    #COL4A6  tandem not annotated in TCGA splice seq
  return = PSI
}

######## A function to rearrange the psi table.
rearrange_psitable = function(psitable)
{
  data = psitable[, 11:ncol(psitable)]
  data1 = as.data.frame(t(data))
  data2 = data.frame(rownames(data1)) #patient ID
  for (i in 1:ncol(data1))
  {
    data_tmp = data.frame(as.numeric(as.character(unname(data1[, i])))) 
    data2 = data.frame(data2, data_tmp)
  }
  colnames(data2) = c("IDs", as.character(psitable[,1]))
  colnames(data2) = paste(colnames(data2), "_psi", sep = "")
  colnames(data2)[1] = "IDs"
  return = data2
}

######## A function to merge the gene level and psi tables by IDs. Currently I have 2 tables
### in the gene expression table IDs look like TCGA-4P-AA8J-01A-11R-A39I-0
### in the spliceseq table IDs look like  TCGA_4P_AA8J or sometimes TCGA_CV_6933_Norm
### I want to have the gne expression like the current spliceseq
### I need a function to bring them all and in the darkness bind them
### I will also add 2 other variables = included_level (psi*gene level) and excluded_level ((1-psi)*gene level)
merge_table = function(exp_table, splice_table)
{
  exp_table[,1] = sub("-", "_", exp_table[,1])
  exp_table[,1] = sub("-", "_", exp_table[,1])
  exp_table[,1] = sub("-", "_", exp_table[,1]) # 3 consecutive "sub" to replace only 3 "-" (no gsub!)
  exp_table[,1] = gsub("-.{1,}", "", exp_table[,1]) # to remove what is behind the previously 4th "-" (now 1st) 
  exp_table[,1] = gsub("11A", "Norm", exp_table[,1]) #norm samples are indexed by 11A in firebrowse, "Norm" in spliceseq
  exp_table[,1] = gsub("_01A", "", exp_table[,1]) ## tumor samples are indexed by 01A in firebrowse, nothing in spliceseq
  exp_table[,1] = gsub("_03A", "", exp_table[,1])
  exp_table[,1] = gsub("_03B", "", exp_table[,1])
  #and now merge
  data_gene = merge(exp_table, splice_table, by = "IDs") 
  ####################
  #prepare columns psi*level, knowing that the genes with psi are included in genes with level 
  indpsi = grep("_psi", colnames(data_gene))
  geneswithpsi = gsub("_psi", "_level", colnames(data_gene)[indpsi])
  indgene = which(colnames(data_gene) %in% geneswithpsi)
  #indpsi and indgene  are the indices of the columns of data_gene with psi and gene level, respectively  
  #but they may not be in the same order and I have to reorder them
  data1 = data_gene[, indpsi]
  order_psi = order(colnames(data1))
  data2 = data1[, order_psi]
  data3 = data_gene[, indgene]
  order_gene = order(colnames(data3))
  data4 = data3[, order_gene]
  data5 = data.frame(data2, data4)
  for (i in 1:ncol(data2))
  {
    data5[, 2*ncol(data2) + i] = data5[, i] * data5[, ncol(data2) + i]
    gene_name_temp = gsub("_psi", "", colnames(data5[i]))
    colnames(data5)[2*ncol(data2) + i] = paste(gene_name_temp, "_included_level", sep = "")
  }
  toto = 2*ncol(data2)+1
  tata = 3*ncol(data2)
  data_gene_1 = data.frame(data_gene, data5[, toto:tata])
  ####################
  # do the same for (1-psi)*level 
  data5 = data.frame(data2, data4)
  for (i in 1:ncol(data2))
  {
    data5[, 2*ncol(data2) + i] = (1 - data5[, i]) * data5[, ncol(data2) + i]
    gene_name_temp = gsub("_psi", "", colnames(data5[i]))
    colnames(data5)[2*ncol(data2) + i] = paste(gene_name_temp, "_excluded_level", sep = "")
  }
  toto = 2*ncol(data2)+1
  tata = 3*ncol(data2)
  data_gene_2 = data.frame(data_gene_1, data5[, toto:tata])
  return = data_gene_2
}

####### A function to plot the distribution of mRNA levels or psi in tumor and normal, adding statistics
plot_gene = function(genes_in_cohort, gene, cohort)
{
  ind = which(colnames(genes_in_cohort) == gene)
  pval = wilcox.test(genes_in_cohort[, ind] ~ genes_in_cohort$type)$p.value
  nNorm = sum(genes_in_cohort$type == "Norm")
  nTum = sum(genes_in_cohort$type == "Tumor")
  ylimmax = max(floor(genes_in_cohort[, ind] - 0.01), na.rm = T)+1 #to get ylim = c(0,1) when psi is computed
  boxplot(genes_in_cohort[, ind] ~ genes_in_cohort$type, ylim = c(0, ylimmax), col = c("#00C65A", "#EFE35C"), main = paste(gene, "\t", cohort, "\n", "Norm n=", nNorm, " Tumor n=", nTum), sub = paste("p (Wilcoxon) = ", pval))
}

############# A function to create a table with clinical data
#requires the clinical data downloaded from firebrowse
# script largely inspired from biostar https://www.biostars.org/p/153013/
format_clinical_data = function(clinical)
{
  # get the columns that contain data we can use: days to death, new tumor event, last day contact to....
  ind_keep <- grep('days_to_new_tumor_event_after_initial_treatment',colnames(clinical))
  # this is a bit tedious, since there are numerous follow ups, let's collapse them together and keep the first value (the higher one) if more than one is available
  new_tum <- as.matrix(clinical[,ind_keep])
  new_tum_collapsed <- c()
  for (i in 1:dim(new_tum)[1]){
    if ( sum ( is.na(new_tum[i,])) < dim(new_tum)[2]){
      m <- min(new_tum[i,],na.rm=T)
      new_tum_collapsed <- c(new_tum_collapsed,m)
    } else {
      new_tum_collapsed <- c(new_tum_collapsed,'NA')
    }
  }
  # do the same to death
  ind_keep <- grep('days_to_death',colnames(clinical)) 
  death <- as.matrix(clinical[,ind_keep])
  death_collapsed <- c()
  for (i in 1:dim(death)[1]){
    if ( sum ( is.na(death[i,])) < dim(death)[2]){
      m <- max(death[i,],na.rm=T)
      death_collapsed <- c(death_collapsed,m)
    } else {
      death_collapsed <- c(death_collapsed,'NA')
    }
  }
  # and days last follow up here we take the most recent which is the max number
  ind_keep <- grep('days_to_last_followup',colnames(clinical))
  fl <- as.matrix(clinical[,ind_keep])
  fl_collapsed <- c()
  for (i in 1:dim(fl)[1]){
    if ( sum (is.na(fl[i,])) < dim(fl)[2]){
      m <- max(fl[i,],na.rm=T)
      fl_collapsed <- c(fl_collapsed,m)
    } else {
      fl_collapsed <- c(fl_collapsed,'NA')
    }
  }
  # and put everything together
  all_clin <- data.frame(new_tum_collapsed,death_collapsed,fl_collapsed)
  colnames(all_clin) <- c('new_tumor_days', 'death_days', 'followUp_days')
  # create vector with time to new tumor containing data to censor for new_tumor
  all_clin$new_time <- c()
  for (i in 1:length(as.numeric(as.character(all_clin$new_tumor_days)))){
    all_clin$new_time[i] <- ifelse ( is.na(as.numeric(as.character(all_clin$new_tumor_days))[i]),
                                     as.numeric(as.character(all_clin$followUp_days))[i],as.numeric(as.character(all_clin$new_tumor_days))[i])
  }
  # create vector time to death containing values to censor for death
  all_clin$new_death <- c()
  for (i in 1:length(as.numeric(as.character(all_clin$death_days)))){
    all_clin$new_death[i] <- ifelse ( is.na(as.numeric(as.character(all_clin$death_days))[i]),
                                      as.numeric(as.character(all_clin$followUp_days))[i],as.numeric(as.character(all_clin$death_days))[i])
  }
  # create vector for death censoring
  all_clin$death_event <- c()
  for (i in 1:length(as.numeric(as.character(all_clin$new_tumor_days)))){
    all_clin$death_event[i] <- ifelse (clinical[i, grep("patient.vital_status", colnames(clinical))]  == 'alive', 0,1)
  }
  #Add patient ID
  all_clin$IDs <- c()
  for (i in 1:length(as.numeric(as.character(all_clin$new_tumor_days)))){
    all_clin$IDs[i] <- clinical[i, grep("patient.bcr_patient_barcode", colnames(clinical))]
      }
  return = all_clin
  }

#####Function to merge the clinical table with the psi+gene level table
### in the clinical table IDs look like tcga-4p-aa8j
### in the gene+psi table IDs look like "TCGA_CV_6933" or "TCGA_CV_6933_Norm"
### of course when I'll merge the 2 tables I'll lose all the data corresponding to the normal tissues (not in the clinical table)
merge_table_all = function(gene_psi_table, clinical_table)
{
  clinical_table$IDs = toupper(clinical_table$IDs) 
  clinical_table$IDs = gsub("-", "_", clinical_table$IDs)
  return = merge(gene_psi_table, clinical_table, by = "IDs")
}

####### A function to identify the best (most discriminative) cut-off between "high" and "low" gene level
# The formula for p value computation is in https://stat.ethz.ch/pipermail/r-help/2007-April/130676.html
# event_rna is a vector with only 0 and 1, where 1 is for the i highest gene values
# This function returns the best cut-off (integer), draws a plot of the pvalue as a function of the cut-off, to check visually the cut-off
#finally it draws the kaplan-meyer curve
kaplan_curve = function(table_values_survival, gene, cohort, excluded_interval = 0.2)
{
  ind = which(colnames(table_values_survival) == gene)
  table_values_survival = table_values_survival[order(table_values_survival[, ind], decreasing = T), ]
  boundary_low = floor(nrow(table_values_survival)*excluded_interval)
  boundary_high = floor(nrow(table_values_survival)*(1-excluded_interval)) #because a cut off that would produce a high imbalance between high and low does not make sense
  toto = matrix(nrow = boundary_high - boundary_low + 1, ncol = 3)
  for (i in boundary_low : boundary_high)
  {
    event_rna = c(rep(1, i), rep(0, nrow(table_values_survival)-i))
    surv =  survdiff(Surv(as.numeric(as.character(table_values_survival$new_death)), table_values_survival$death_event) ~ event_rna)
    pval = 1 - pchisq(surv$chisq, length(surv$n) - 1)
    toto[i - boundary_low +1 ,1] = i
    toto[i - boundary_low +1 ,2] = pval 
  }
  toto[,3] = -log10(toto[,2])
  plot(toto[,3] ~ toto[,1])  #-log10(p value) vs cut off position
  oco1 = which.max(toto[ ,3]) #optimal cut off
  oco = oco1 + boundary_low - 1 #to get the index in th whole table, not only in 0.2-0.8
  event_rna = c(rep(0, oco), rep(1, nrow(table_values_survival)-oco))
  surv =  survdiff(Surv(as.numeric(as.character(table_values_survival$new_death)), table_values_survival$death_event) ~ event_rna)
  pval = 1 - pchisq(surv$chisq, length(surv$n) - 1)
  toto = survfit(Surv(as.numeric(as.character(table_values_survival$new_death)), table_values_survival$death_event) ~ event_rna)
  plot(survfit(Surv(as.numeric(as.character(table_values_survival$new_death)), table_values_survival$death_event) ~ event_rna), mark.time = T, main = paste(gene, cohort, sep = "  "), xlab = "days", col = c(1, 4), frame = F, lwd = 2)
  mtext(paste("p (log rank) = ", pval), side = 3)
  text(yprint, 1, paste("High, n=", toto$n[1], " median=", unname(summary(toto)$table[, "median"][1])))
  text(yprint, 0.9, paste("Low, n=", toto$n[2], " median=", unname(summary(toto)$table[, "median"][2])), col = "blue")
  return = c(unname(summary(toto)$table[, "median"][1]), unname(summary(toto)$table[, "median"][2]), toto$n[1], toto$n[2], pval)
}

#### A function to draw survival curves combining CELF1 and ELAVL1 level
#I want to take on the one hand all the cancers with low CELF1 AND ELAVL1 levels, and on the other hand the other ones (high CELF1 OR ELAVLA)
# "high" and "low" are defined with cut-offs optimised individually for CELF1 and ELAVL1 as above
#So I prepare a vector (event_rna) with "1" for the bottom values of CELF1, another vector with "1" for the bottom values of CELF1, and by multiplying the vectors I have "1" for bottom values of both
#this function also returns a data.frame enriched with a "elav_celf_co" column equal to 0 (high CELF or ELAV) or 1 (low CELF and ELAV)
kaplan_curve_CE = function(table_values_survival, excluded_interval = 0.2)
{
  boundary_low = floor(nrow(table_values_survival)*excluded_interval)
  boundary_high = floor(nrow(table_values_survival)*(1-excluded_interval)) 
  #start with defining the cutoff for CELF1
  data = table_values_survival
  ind = which(colnames(data) == "CELF1_level")
  data = data[order(data[, ind], decreasing = T), ]
  toto = matrix(nrow = boundary_high - boundary_low + 1, ncol = 3)
  for (i in boundary_low : boundary_high)
  {
    event_rna = c(rep(1, i), rep(0, nrow(data)-i))
    surv =  survdiff(Surv(as.numeric(as.character(data$new_death)), data$death_event) ~ event_rna)
    pval = 1 - pchisq(surv$chisq, length(surv$n) - 1)
    toto[i - boundary_low +1 ,1] = i
    toto[i - boundary_low +1 ,2] = pval 
  }
  toto[,3] = -log10(toto[,2])
  oco1 = which.max(toto[ ,3]) #optimal cut off
  oco_CELF = oco1 + boundary_low - 1 #to get the index in th whole table, not only in 0.2-0.8
  #now the cutoff for ELAVVLF1
  ind = which(colnames(data) == "ELAVL1_level")
  data = data[order(data[, ind], decreasing = T), ]
  toto = matrix(nrow = boundary_high - boundary_low + 1, ncol = 3)
  for (i in boundary_low : boundary_high)
    {
    event_rna = c(rep(1, i), rep(0, nrow(data)-i))
    surv =  survdiff(Surv(as.numeric(as.character(data$new_death)), data$death_event) ~ event_rna)
    pval = 1 - pchisq(surv$chisq, length(surv$n) - 1)
    toto[i - boundary_low +1 ,1] = i
    toto[i - boundary_low +1 ,2] = pval  
  }
  toto[,3] = -log10(toto[,2])
  oco1 = which.max(toto[ ,3]) #optimal cut off
  oco_ELAV = oco1 + boundary_low - 1 #to get the index in th whole table, not only in 0.1-0.9
  ## now we combine both
  data = data[order(data$ELAVL1_level, decreasing=T), ]
  data$elav_co = c(rep(0, oco_ELAV), rep(1, nrow(data)-oco_ELAV))
  data = data[order(data$CELF1_level, decreasing=T), ]
  data$celf_co = c(rep(0, oco_CELF), rep(1, nrow(data)-oco_CELF))
  data$elav_celf_co = data$elav_co * data$celf_co #the value is 1 only if both elav_co and celf_co have 1, ie low CELF AND ELAV
  surv =  survdiff(Surv(as.numeric(as.character(data$new_death)), data$death_event) ~ data$elav_celf_co)
  pval = 1 - pchisq(surv$chisq, length(surv$n) - 1)
  toto = survfit(Surv(as.numeric(as.character(data$new_death)), data$death_event) ~ data$elav_celf_co )
  plot(survfit(Surv(as.numeric(as.character(data$new_death)), data$death_event) ~ data$elav_celf_co), mark.time = T, xlab = "days", main = paste("CELF and ELAV ", cohort), col = c(1,4), frame = F, lwd = 2)
  mtext(paste("p (log rank) = ", pval), side = 3)
  text(yprint, 0.9, paste("High C or E, n=", toto$n[1], " median=", unname(summary(toto)$table[, "median"][1])))
  text(yprint, 0.8, paste("Low C and E, n=", toto$n[2], " median=", unname(summary(toto)$table[, "median"][2])), col = "blue")
  data$elav_celf_co[data$elav_celf_co == "1"] = "Low_CELF_and_ELAV"
  data$elav_celf_co[data$elav_celf_co == "0"] = "High_CELF_or_ELAV"
  data$elav_celf_co = as.factor(data$elav_celf_co)
  return = list(data, c(unname(summary(toto)$table[, "median"][1]), unname(summary(toto)$table[, "median"][2]), toto$n[1], toto$n[2], pval))
}

#### A function to plot a gene level as a function of low C and E vs high C or E, adding statistics
plot_gene_CE = function(genes_in_cohort, gene, cohort)
{
  ind = which(colnames(genes_in_cohort) == gene)
  pval = wilcox.test(genes_in_cohort[, ind] ~ genes_in_cohort$elav_celf_co)$p.value
  nLow = sum(genes_in_cohort$elav_celf_co == "Low_CELF_and_ELAV")
  nHigh = sum(genes_in_cohort$elav_celf_co == "High_CELF_or_ELAV")
  deltapsi = median(genes_in_cohort[genes_in_cohort$elav_celf_co == "High_CELF_or_ELAV", ind], na.rm = T) -  median(genes_in_cohort[genes_in_cohort$elav_celf_co == "Low_CELF_and_ELAV", ind], na.rm = T)
  ylimmax = max(floor(genes_in_cohort[, ind] - 0.01), na.rm = T)+1 #to get ylim = c(0,1) when psi is computed
  boxplot(genes_in_cohort[, ind] ~ genes_in_cohort$elav_celf_co, ylim = c(0, ylimmax), col = c("#00C65A", "#EFE35C"), main = paste(gene, "\t", cohort, "\n", "High CELF or ELAV n=", nHigh, " Low CELF and ELAV n=", nLow), cex.main = 1, sub = paste("p (Wilcoxon) = ", pval, "\n", "DeltaPSI = ", deltapsi))
  return = c(pval, nLow, nHigh, deltapsi)
}

#### A function that returns the the PSI values depending on CELF1 and ELAV levels being in the top or bottom XX% (quant)
#can also return a boxplot (commented line)
plot_gene_CE_quantile = function(genes_in_cohort, gene, cohort, quant = 0.5)
{
  data = genes_in_cohort
  ind_gene = which(colnames(data) == gene)
  thresholdHigh =  floor(nrow(data) * quant) # the number of patients to be considered as expressing the highest levels of C or E
  thresholdLow =  floor(nrow(data) * (1-quant))+1 #same for bottom expression
  #First sort with respect to top or bottom CELF1 level
  ind = which(colnames(data) == "CELF1_level")
  data = data[order(data[, ind], decreasing = T), ]
  data$CELF_quant = 10 #first put 10 everywhere
  data$CELF_quant[1 : thresholdHigh] = 1 #1 for the top quant of CELF1 levels
  data$CELF_quant[thresholdLow : nrow(data)] = 0  #0 for the bottom quant of CELF1 levels
  #Then sort with respect to top or bottom ELAVL1 level
  ind = which(colnames(data) == "ELAVL1_level")
  data = data[order(data[, ind], decreasing = T), ]
  data$ELAV_quant = 10 #first put 10 everywhere
  data$ELAV_quant[1 : thresholdHigh] = 1 #1 for the top quant of CELF1 levels
  data$ELAV_quant[thresholdLow : nrow(data)] = 0  #0 for the bottom quant of CELF1 levels
  #combine both
  data$CELFELAV =  data$CELF_quant + data$ELAV_quant #0 if both C and E in the bottom quant, 2 if both C and E in the top quant, 1 or 10 or 11 or 20 otherwise
  # low C and E vs high C and E, ie data$CELFELAV == 0 vs data$CELFELAV == 2
  data = data[data$CELFELAV == 0 |  data$CELFELAV == 2, ]
  pval = wilcox.test(data[, ind_gene] ~ data$CELFELAV)$p.value
  nLow = sum(data$CELFELAV == 0)
  nHigh = sum(data$CELFELAV == 2)
  medlow = median(data[, ind_gene][data$CELFELAV == 0], na.rm = T)
  medhigh = median(data[, ind_gene][data$CELFELAV == 2], na.rm = T)
  boxplot(data[, ind_gene] ~ data$CELFELAV, col = c("#00C65A", "#EFE35C"), main = paste(gene, "\t", cohort, "\n", "High CELF and ELAV n=", nHigh, " Low CELF and ELAV n=", nLow), cex.main = 1, sub = paste("p (Wilcoxon) = ", pval), names = c("low", "high"))
  return = c(medhigh, nHigh, medlow, nLow, pval)
 }

######################
####### Read the TCGA tables, rearrange them, merge them into a single table
library(survival)
cohort = "LUSC"
wd = paste("~/Luc_travail/recherche_divers/TCGA_and_co/", cohort, sep = "")
setwd(wd)
gene_set_level = c(as.character(read.table("diff_probes_FDR<0.1", h=T)$GeneName), "CELF1", "ELAVL1")
gene_set_psi = as.character(read.table("diff_probes_FDR<0.1", h=T)$GeneName)
#Since I need 3 big tables I successively read them, work with them to reduce their size and remove them from the environment
## first check if I have useful clinical data before proceeding with gene elvels etc
dataclin = paste(cohort, ".merged_only_clinical_clin_format.txt", sep = "")
data_clinical = read.csv(dataclin, header=TRUE, row.names = 1, sep="\t", fill = T)
#data_clinical = read.csv("CHOL.clin.merged.txt", header=TRUE, row.names = 1, sep="\t", fill = T)
data6 = t(data_clinical)
data7 = format_clinical_data(clinical = data6); head(data7)
rm(data_clinical)
datagene = paste(cohort, ".rnaseqv2__illuminahiseq_rnaseqv2__unc_edu__Level_3__RSEM_genes_normalized__data.data.txt", sep = "")
data_gene = read.table(datagene, h=F, sep = "\t")
#data_gene = read.table("COAD.rnaseqv2__illuminaga_rnaseqv2__unc_edu__Level_3__RSEM_genes_normalized__data.data.txt", h=F, sep = "\t")
#data_gene = read.table("READ.rnaseqv2__illuminaga_rnaseqv2__unc_edu__Level_3__RSEM_genes_normalized__data.data.txt", h=F, sep = "\t")
#data_gene = read.table("UCEC.rnaseqv2__illuminaga_rnaseqv2__unc_edu__Level_3__RSEM_genes_normalized__data.data.txt", h=F, sep = "\t")
data1 = gene_levels(data_genexp = data_gene, gene_set = gene_set_level)
data2 = add_type(data_gene = data1); head(data2)
rm(data_gene)
data_PSI = read.table("PSI_download_allgenes.txt", h=T)
data3 = PSI_goi(PSI_download = data_PSI, gene_set = gene_set_psi)
data4 = clean_PSI_selected(PSI_CELFELAV = data3)
data5 = rearrange_psitable(psitable = data4); head(data5)
rm(data_PSI)
# Prepare a general table (merge of clinical data, gene levels, and PSI)
data_level_psi = merge_table(exp_table = data2, splice_table = data5)
data_all = merge_table_all(gene_psi_table = data_level_psi, clinical_table = data7)
head(data_all)

#draw the survival curves and prepare table for TCGA pvalues
yprint = 3000
tmp_celf = kaplan_curve(table_values_survival = data_all, gene = "CELF1_level", cohort = cohort)
tmp_elav = kaplan_curve(table_values_survival = data_all, gene = "ELAVL1_level", cohort = cohort)
list_data_CE = kaplan_curve_CE(table_values_survival = data_all, excluded_interval = 0.1)
tmp_CE = list_data_CE[[2]] #a vector with survival medians, n and p value
tmp_CE[3] / (tmp_CE[3] + tmp_CE[4]) #to compare the numbers in high and low
tmp = c(tmp_celf, tmp_elav, tmp_CE)
#names_tmp = c("celf_median_high", "celf_median_low", "celf_nHigh", "celf_nLow", "celf_pvalue", "elav_median_high", "elav_median_low", "elav_nHigh", "elav_nLow", "elav_pvalue", "CE_median_high", "CE_median_low", "CE_nHigh", "CE_nLow", "CE_pvalue")
#table_TCGA = data.frame(names_tmp, tmp) #for the first round (first cohort)
#then I do the same for each cohort and I paste the result of each cohort with the previous ones
table_TCGA = data.frame(table_TCGA, tmp) #for the subsequent rounds
colnames(table_TCGA)[colnames(table_TCGA) == "tmp"] = cohort
#once the job is done for all the cohorts I have to rearrange the data.frame
table_TCGA_1 = t(table_TCGA[ , 2:ncol(table_TCGA)])
colnames(table_TCGA_1) = table_TCGA$names_tmp
# write.table(table_TCGA_1, "table_TCGA") #at the end of the process

### Barplot the pvalues of survival curves, low vs high CELF, ELAV, or both
setwd("~/Luc_travail/Papiers/CEFL1-ELAVL1/manuscrit_aut18/R_script_and_results")
#The file "table_TCGA" contains the cohort name, with the pvalues of kaplan meier curves
data = read.table(file = "table_TCGA", h=T)
head(data)
data$Cohort_name = rownames(data)
data = data[order(data$Cohort_name, decreasing = T), ]  # to have the 1st cohorts in alphabetical order at the top of the graph
#remove the cohorts where CELF and ELAV have different effects (based on visual inspection of curves)
cohort_remove = c("THYM", "TGCT", "SKCM", "READ", "OV", "LUAD", "LIHC", "KIRP", "KICH", "GBM", "CESC")
data = data[!(data$Cohort_name %in% cohort_remove),]
#remove the cohorts with ridiculous pvalues for both C and E
data = data[data$celf_pvalue < 0.1 | data$elav_pvalue < 0.1,]
#remove the cohorts with one protein that alone provides a good stratification
data = data[data$celf_pvalue > 0.01 & data$elav_pvalue > 0.01,]
#and now the graph
mat = cbind(-log10(data$celf_pvalue), -log10(data$elav_pvalue), -log10(data$CE_pvalue)) 
dimnames(mat) = list(data$Cohort_name, c("CELF", "ELAV", "CE"))
barplot(t(mat), beside = T, horiz = T, cex.names = 0.6, las = 1, col = c("green", "blue", "orange"), legend.text = T, space = c(0,2), xlab = "-log10(pvalue)")
abline(v=1, lty=2)
abline(v=3, lty=2)

### Compute CD44 PSI in high C+E vs low C+E, and plot the values adding statistics
#don't forget to re-prepare data_all for the cohorts of interest
plot_gene_CE_quantile(genes_in_cohort = data_all, gene = "CD44_psi", cohort = cohort, quant = 0.5)
## Draw KM curves for CD44_PSI
yprint = 2000
kaplan_curve(table_values_survival = data_all, gene = "CD44_psi", cohort = cohort)
